# Joint representation and visualization of derailed cell states with Decipher

**DOI:** 10.1101/2023.11.11.566719

**Authors:** Achille Nazaret, Joy Linyue Fan, Vincent-Philippe Lavallée, Cassandra Burdziak, Andrew E. Cornish, Vaidotas Kiseliovas, Robert L. Bowman, Ignas Masilionis, Jaeyoung Chun, Shira E. Eisman, James Wang, Justin Hong, Lingting Shi, Ross L. Levine, Linas Mazutis, David Blei, Dana Pe’er, Elham Azizi

**Affiliations:** Department of Computer Science, Columbia University, New York, NY 10027, USA; Irving Institute for Cancer Dynamics, Columbia University, New York, NY 10027, USA; Department of Biomedical Engineering, Columbia University, New York, NY 10027, USA; Computational and Systems Biology Program, Sloan Kettering Institute, Memorial Sloan Kettering Cancer Center, New York, NY 10065, USA; Centre Hospitalier Universitaire Sainte-Justine Research Center, Montréal, QC, Canada; Department of Pediatrics, Université de Montréal, Montréal, QC, Canada; Immunology Program, Memorial Sloan Kettering Cancer Center, New York, NY 10065, USA; Department of Medicine, Memorial Sloan Kettering Cancer Center, New York, NY 10065, USA; Alan and Sandra Gerry Metastasis and Tumor Ecosystems Center, Memorial Sloan Kettering Cancer Center, New York, NY 10065, USA; Human Oncology and Pathogenesis Program, Memorial Sloan Kettering Cancer Center, New York, NY 10065, USA; Department of Cancer Biology, University of Pennsylvania, Philadelphia, PA 19104, USA; Institute of Biotechnology Vilnius University, Life Sciences Centre, Vilnius 02158, Lithuania; Department of Statistics, Columbia University, New York, NY 10027, USA; Data Science Institute, Columbia University, New York, NY 10027, USA; Howard Hughes Medical Institute, Memorial Sloan Kettering Cancer Center, New York 10027, NY 10065, USA; Herbert Irving Comprehensive Cancer Center, Columbia University, New York, NY 10032, USA

**Author notes:** equal contribution (Achille Nazaret, Joy Linyue Fan, Vincent-Philippe Lavallée).

## Abstract

Biological insights often depend on comparing conditions such as disease and health, yet we lack effective computational tools for integrating single-cell genomics data across conditions or characterizing transitions from normal to deviant cell states. Here, we present Decipher, a deep generative model that characterizes derailed cell-state trajectories. Decipher jointly models and visualizes gene expression and cell state from normal and perturbed single-cell RNA-seq data, revealing shared and disrupted dynamics. We demonstrate its superior performance across diverse contexts, including in pancreatitis with oncogene mutation, acute myeloid leukemia, and gastric cancer.

## BACKGROUND

Single-cell genomic technologies have enabled the detailed characterization of cellular states in healthy and disease contexts, including cancer [1–5], inflammatory bowel disease [6,7], and COVID-19 [8–10]. Single-cell RNA sequencing (scRNA-seq) of single tissue snapshots can characterize cell-state transitions by applying pseudotime inference approaches that leverage the asynchrony of cell states in biological samples [11–15]. In particular, reconstructing how cells derail from normal to diseased states along a pseudotime axis promises to improve our knowledge of early disease stages, identify drivers of this derailment, and inform early detection and prevention strategies.

A prime example of derailed development occurs in acute myeloid leukemia (AML), a lethal cancer of the hematopoietic system. In AML, bone marrow hematopoietic stem and progenitor cells (HSPCs) acquire genetic and epigenetic abnormalities, leading to the accumulation of HSPC-like leukemic cells called “blasts” that fail to differentiate terminally. The origin of pre-leukemic and leukemic stem cell states in AML [5,16] remains poorly characterized, which makes it difficult to target these cells and prevent disease recurrence [17–19]. Importantly, we do not know how specific mutations lead to distinct disease trajectories. These trajectories differ across patients[20] and can initiate from healthy states, or pre-malignant and early malignant states such as clonal hematopoiesis and myelodysplastic syndromes [21–25]. Numerous other contexts, including disorders of embryonic development, neurodegenerative diseases, and T-cell exhaustion[26–29], require the accurate reconstruction of aberrant trajectories to understand their mechanisms.

However, existing methods often fail to faithfully reconstruct the order of events; linear approaches, such as principal component analysis (PCA), cannot capture biological complexity, while alternatives, such as neural networks, typically fail to represent the underlying biology and can mix the ordering of cell states. There is an urgent need for methods to accurately reconstruct the order of transcriptional events, precisely align trajectories, and compare disparate conditions, such as healthy to disease and control to genetic or chemical perturbation.

To compare trajectories, it is critical to obtain a faithful joint embedding and accurately visualize the cellular relationships it represents. Embedding multiple samples, especially from heterogeneous cancers, is sensitive to minor differences in gene programs between samples, such that they typically fail to co-embed in a biologically meaningful way. Existing integration methods[30–37] are primarily designed for batch correction; they assume that samples share similar cell states and attempt to eliminate differences—including genuine biological differences, particularly for continuous and diverging trajectories—as technical effects. Moreover, most approaches compress information from thousands of genes into 10–50 factors that are independent, thereby neglecting dependencies between related biological processes (ignoring, for example, that divergent differentiation trajectories are related). The resulting latent spaces help annotate discrete cell states but often do not preserve gene-gene relationships and the order of cell states [38]. Furthermore, data is usually visualized by projecting latent embeddings onto two dimensions [39]–[40], which can distort topology and obscure functional relationships [41]. These limitations highlight the need for approaches that address interpretability, preserve global geometry in the latent space and enable visualization to better model trajectories perturbed by mutation, genetic manipulation, drugs, or disease.

In this work, we present Decipher (deep characterization of phenotypic derailment), an interpretable deep generative model for the simultaneous integration and visualization of single-cell data from disparate conditions. Decipher is a hierarchical model that learns two representations for each cell from observed expression: a low-dimensional state (in an ‘intermediate’ latent space of roughly ten dimensions, similar to existing methods [37,42]), as well as a two-dimensional representation (‘top’ latent space) for visualization. Several design features allow this unifying model to characterize continuous trajectories more accurately: 1) it connects gene expression and the latent spaces with simple linear or one-layer neural network transformations to limit distortion, 2) the stacking of two latent spaces over gene expression space enables flexible capture of nonlinear mechanisms, despite the use of simple transformations, 3) it learns the dependency structure of cell-state latent factors with the top latent space embedding (unlike other methods, which assume that latent factors are independent), enabling the discovery of both shared and unique biological mechanisms from sparse trajectories, and 4) the 2D top latent space provides a direct visualization of the geometry learned by the model.

The hierarchical structure of Decipher allows for a comprehensive understanding of the relationships between gene expression, cell states, and their visual representation. We show that it is the only approach that preserves cell-state organization and continuity in synthetic data and demonstrate its substantial advantage for deriving insight from three disease contexts of increasing complexity—published data from a pancreatic ductal adenocarcinoma (PDAC) mouse model with an oncogenic mutation, new data from heterogeneous AML patient specimens, and a published patient cohort spanning two subtypes of gastric cancer.

## RESULTS

### The Decipher method

Aligning trajectories from normal and perturbed contexts requires a joint representation that preserves the topology and order of cells along both trajectories without forcing artificial overlap. Decipher’s key assumption is that perturbed trajectories maintain shared transcriptional programs with normal trajectories for common processes, such as cell maturation. To create a joint representation that captures both biological differences and shared mechanisms, Decipher employs a hierarchical model featuring two levels of latent representations, each with its own encoder and decoder networks. This unique architecture allows for correlation between some latent factors, enabling the identification of shared gene programs that would be missed under the standard requirement for independence. Decipher uses simple linear transformations and single-layer neural networks to connect all representations within a unified probabilistic framework, making it sufficiently flexible to learn nonlinear mechanisms while imposing a rigid inductive bias that prevents arbitrary distortion of the global geometry.

To enable correlated latent factors, Decipher extends the successful single-cell variational auto-encoder (VAE) architecture [37,42–44] into a deeper generative model inspired by the deep exponential family [45,46]. In Decipher, each cell has not one but two complementary latent representations (**Fig. 1a,b**). First, we embed cells in a two-dimensional representation that encodes global cell-state dynamics. We refer to this high-level embedding as the *Decipher space* and its two dimensions as *Decipher components*. Decipher components represent the dominant axes of variation in the data, typically progression (maturation) and derailment (degree of deviation from a normal process, e.g. in disease). Then, we generate a higher-dimensional representation conditional on the two Decipher components, designed to capture more refined cell-state information. We call the space of refined representations the *latent space* and refer to its dimensions as *latent factors*. The latent space is similar in principle to previous VAEs [37,43,44], except that the design of Decipher, which conditions the latent space on the Decipher components, enables dependencies between latent factors. Finally, Decipher generates the denoised gene expression of each cell from its inferred cell state (**Fig. 1b** and Methods).

**Figure 1.**
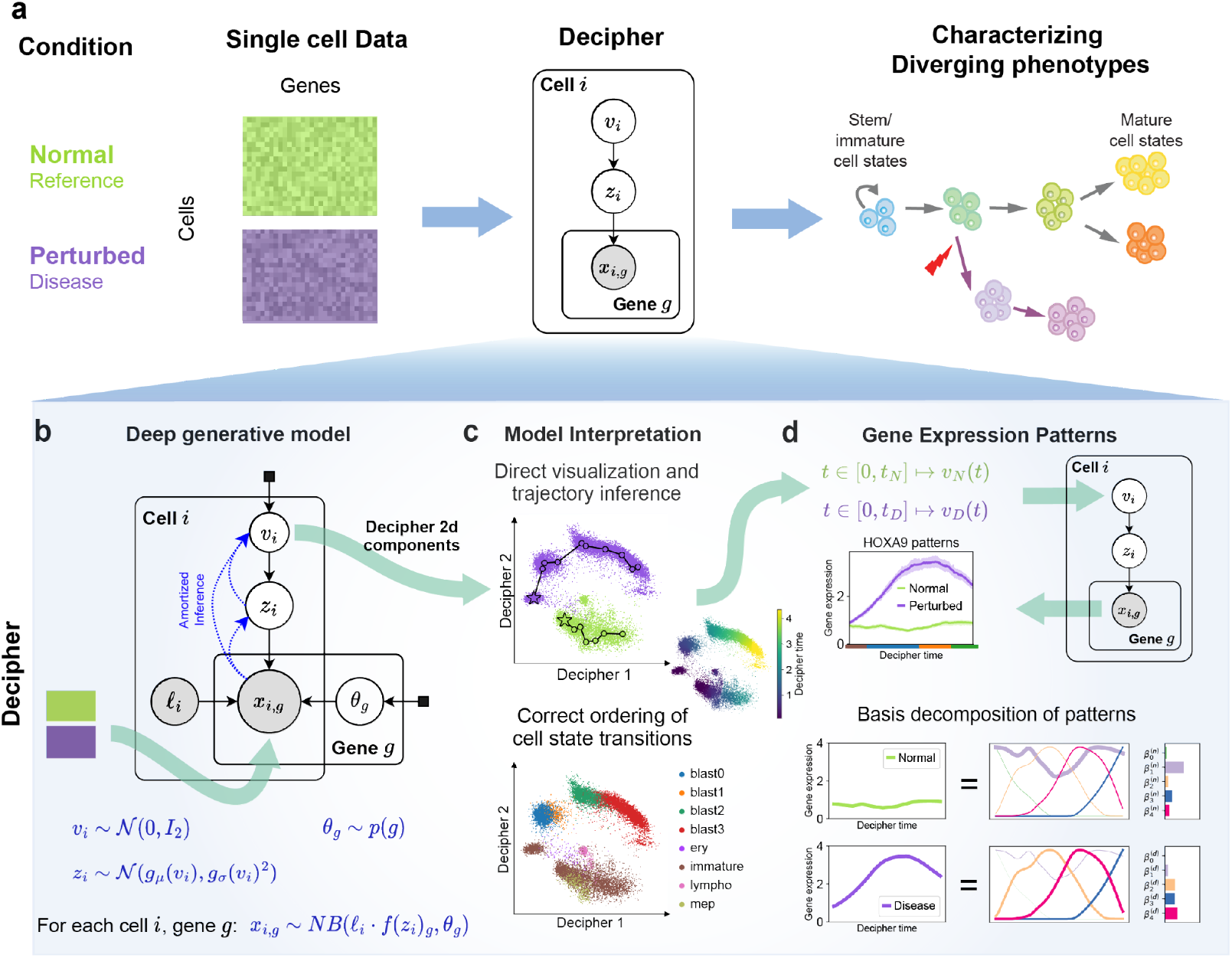
Overview of the Decipher framework. **(a)** Decipher accepts two single-cell datasets from a normal (reference) and a perturbed (e.g., disease) condition as input, which its probabilistic model learns to represent in a hierarchical shared latent space without removing biological differences. The latent space reveals shared cell-state transitions and characterizes diverging phenotypes. **(b)** The Decipher generative model has three levels of cell representations (distributions shown at bottom): 2D Decipher components *v*, latent factors *z*, and gene expression *x*. Decipher components summarize heterogeneous cell states and are used to directly visualize the latent space. **(c)** Example of Decipher space visualization, colored by the dataset of origin (normal or disease) and by manually annotated cell states. Two distinct trajectories (lines and circles; stars depict start) are computed in the Decipher space. **(d)** Gene expression patterns are computed along each trajectory using Decipher’s mapping from *v* to *x* and then decomposing into representative patterns (basis). The corresponding weights are used to compare patterns between the two contexts to pinpoint disrupted and conserved genes.

This three-step generative process interpolates between different degrees of non-linearity and generates latent factors whose dependencies are shaped by high-level Decipher components, offering major advantages for interpretation. Cell representations can be visualized in 2D directly from Decipher components, eliminating the need for further dimensionality reduction with methods such as t-distributed stochastic neighbor embedding (tSNE) or uniform manifold approximation and projection (UMAP), whose usage is a subject of debate [47] (**Fig. 1c**). Within the Decipher space, derailed trajectories can be constructed along a joint pseudotime that we call *Decipher time*.

In addition, Decipher is formulated to give explicit mapping functions between gene space and Decipher space, enabling a straightforward reconstruction of gene expression anywhere in Decipher space (Methods) and the imputation of gene trends along the entire trajectory (**Fig. 1d**). This is particularly useful for determining gene expression levels in sparse locations of the Decipher space with few observed cells. It also enables the reciprocal computation of Decipher components (and straightforward visualization) for any cell with measured gene expression. In contrast, there is no explicit mapping between UMAP space and gene expression space. Decipher offers a unique framework for dimensionality reduction, 2D visualization, trajectory alignment, and characterization of cell state transitions.

### Decipher preserves sparse simulated trajectories

To benchmark Decipher against alternative methods, we simulated ground truth continuous cell-state trajectories by randomly sampling two-dimensional vectors (representing cell states) along a forked trajectory containing regions of low (0 to 10%) sampling density (**Fig. 2a**). The sparsely sampled regions reflect realistic variation in data collected from snapshots of a stepwise differentiation process[14]. We transformed ground truth cell-states into gene expression using random neural networks, similar to scVI’s generative process (Methods), then visualized the data using popular dimensionality approaches (force-directed layout[40], UMAP[40,48], PHATE[49], scVI[37]), and measured how well they recover the true organization of cell states.

**Figure 2.**
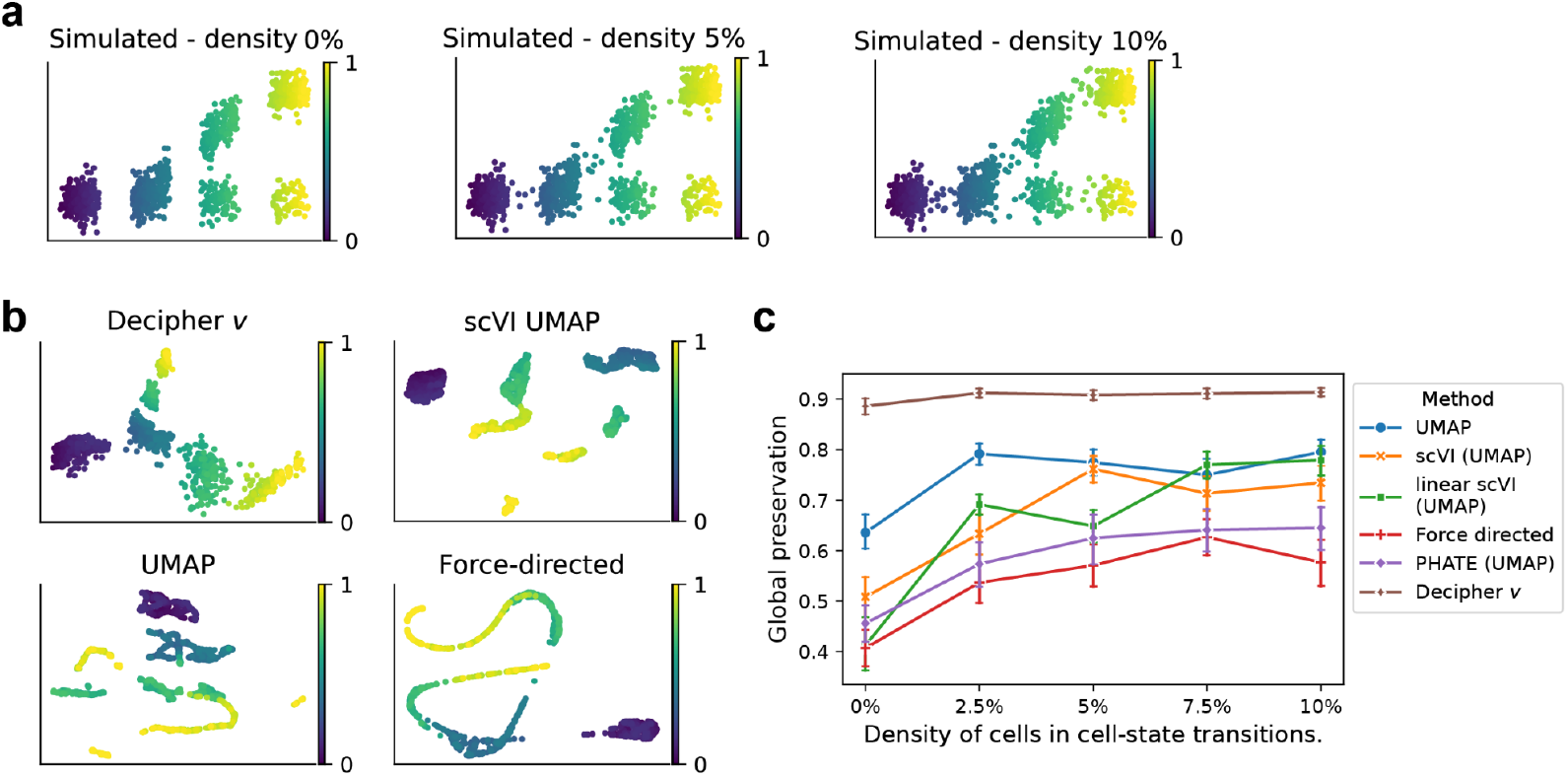
Comparison of methods on simulated data. **(a)** Simulated cell states along diverging trajectories with downsampled (0% to 10% cell density) cell-state transitions. Color gradient represents ground-truth simulated pseudotime. **(b)** Latent spaces learned by different methods on ground truth data with cell-state transition density of 0%. Only Decipher preserves the global order of cell states. **(c)** Global preservation of five independent random datasets across a range of cell-state transition densities, by method (Methods; 1 indicates best preservation). Error bars, s.d. Decipher best preserves the global order of cell states for all cell-state transition densities.

Only Decipher produced visualizations that reflect the two trajectories in the correct order (**Fig. 2b**). Errors made by the other tested methods, such as the close proximity of initiating and terminal cells in the scVI latent space visualized with UMAP, are caused by low cell densities in transitional regions[50]. Although cell-state transitions are common and important in biology, current methods are only designed to preserve distances in locally continuous data, and thus lose the global geometry of cell states. We quantitatively evaluated the latent representation using a global preservation metric[41], which measures the accuracy of cell-state ordering by first computing a nearest-neighbor graph on ground-truth data clustered into cell states, then determining whether neighbors are retained in the learned visualization (Methods). Decipher space exhibits much greater global preservation than the other methods across a realistic range[51] of transitional cell densities, with the most pronounced improvement in lowly sampled regimes (**Fig. 2c**).

### Decipher improves the interpretation of oncogenic trajectories

We next evaluated whether Decipher can characterize the impact of oncogenic *Kras* mutation on pancreas regeneration in mice [52]. Following injury, wild-type epithelial cells undergo physiological metaplasia and regeneration, whereas *Kras*-mutated cells enter a premalignant state that begins expressing oncogenic programs and presages cancer [53] (**Fig. 3a**). Decipher’s 2D space successfully separates wild type (‘normal’) from mutant conditions and organizes cells into three smooth visual trajectories corresponding to two normal conditions and a *Kras-*mutated condition (**Fig. 3b**).

**Figure 3.**
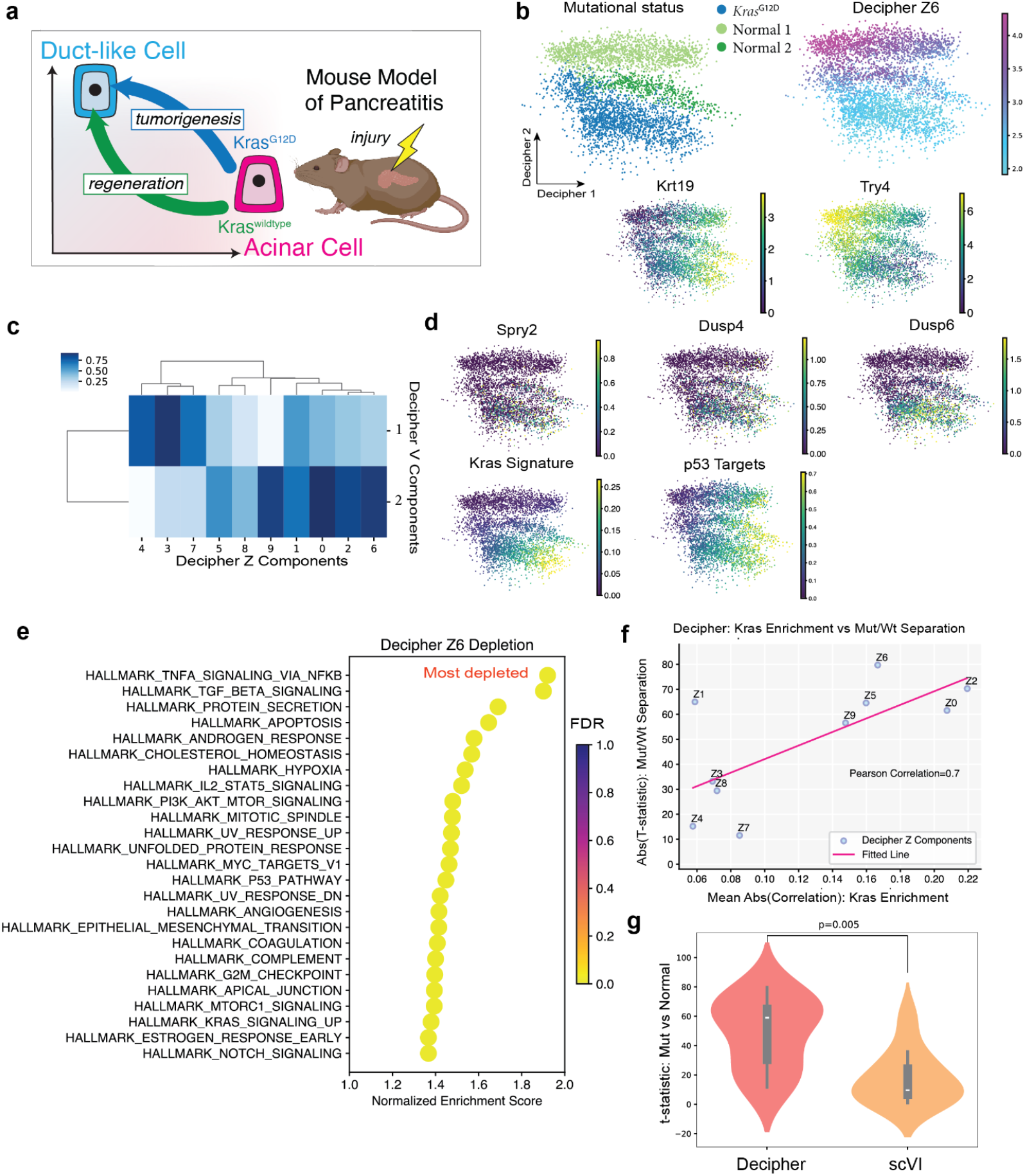
Decipher delineates the impact of *Kras* mutation on pancreatic regeneration. **(a)** Model of pancreatitis in mice. Injury drives acinar cells to ductal-like cell states, aiding regeneration in wild type but promoting tumorigenesis in an oncogenic *Kras* background. **(b)** Decipher 2D space colored by *Kras* mutation status, latent factor z6 loading, or acinar (*Try4*) or ductal (*Krt19*) marker expression. **(c)** Pearson correlation between Decipher components (v) and latent factors (z). **(d)** Decipher 2D space colored by expression of the *Kras* mutational signature, p53 targets, and *Kras* targets (*Dusp4, Dusp6, Spry2*). **(e)** Pathways depleted in Decipher z6. **(f)** Correlation between the absolute value of the t-statistic quantifying the distance between *Kras*-mutant and normal cells in each latent factor, and the mean absolute value of each factor’s enrichment in the bulk *Kras* mutational signature. **(g)** *t* statistic comparing the distributions of each latent component in *Kras*-mutated (*Kras*^G12D^) versus normal cells for Decipher and scVI.

Importantly, Decipher 1 is able to capture the well-known process of acinar-to-ductal metaplasia (ADM), which appears as a smooth progression from acinar (*Try4+*) to ductal (*Krt19+*) cells (**Figs. 3b** and **S1a;** Methods). ADM is a normal regenerative response to injury that occurs in both healthy and disease systems[54],[55], highlighting Decipher’s ability to model shared cellular processes. At the same time, Decipher 2 delineates the derailed trajectory to *Kras*-induced premalignant states, separating trajectories by the degree of deviation from normal, while maintaining their alignment to the shared ADM process. Decipher correctly identifies that one normal condition is more similar to the *Kras-*mutated condition, as supported by the shared expression of important regulators such as *AP1*, likely induced by stress which also occurs during normal regeneration(**Fig. S1b**). By faithfully representing the global geometry of the data, Decipher thus generates highly interpretable 2D components (v).

The latent factors (z) learned by Decipher offer further insight into the derailment, by distinguishing key cell populations (**Fig. S1c**) and revealing which genes inform these distinctions. We identified the top factor separating *Kras*-mutant and normal cells, and found that this factor (z6) highlights the *Kras*-mutated population and is strongly driven by Decipher 2 (**Fig. 3b,c** and Methods). To identify genes associated with z6, we computed the correlation between the expression of genes across all cells and latent factor weights. Notably, *Kras* target genes *Dusp6, Dusp4, Spry2* and *Spry4* were ranked significantly higher, by correlation with negative z6, than the ranking distribution of all genes (*p* = 0.006, Wilcoxon rank sum test; **Fig. 3d** and **Table 1**). Gene set enrichment analysis on factor z6 using these correlation-based gene rankings identified 42 significantly enriched pathways (FDR *q* < 0.25; **Table 2**), including TNF, TGFB, MYC and p53 pathways associated with tumorigenesis (**Fig. 3e**). Finally, a *Kras* mutational signature derived from bulk data [52] increases along the Decipher 1 axis and is only enriched in the *Kras* mutated population of cells, while *p53* targets [56] increase along all three trajectories (**Fig. 3d**), reflecting p53’s intact status at this premalignant stage. These results support Decipher’s ability to dissect premalignant states and clearly illustrate the derailment from normal regeneration caused by a single oncogenic mutation.

Notably, the z factors that most effectively distinguish *Kras-*mutated from wild-type cells (as determined by *t*-test) are also the most correlated with the bulk *Kras* mutational signature (**Fig. 3f**), demonstrating that Decipher decomposes the effects of regeneration and *Kras* mutation in this process. We found that the latent space of scVI, though also VAE-based, is less effective at separating normal and *Kras*-mutated cells (**Figs. 3g** and **S1d,e**). More importantly, the *Kras* mutational signature is very poorly correlated with mutational state separation across scVI latent factors, and gene set enrichment analysis of these factors also fails to find many relevant pathways (**Fig. S1f,g**).

Decipher can recover many more pathways associated with the *Kras*-deviated trajectory since it allows dependencies between factors, which helps identify shared and distinct features of each sample (**Fig. 3c**). By simultaneously capturing the shared physiological metaplasia (via Decipher 1) and the distinct oncogenic derailment (via Decipher 2 and z6), Decipher provides a robust framework for interpreting both normal and disease trajectories. This dual capability for modeling and visualizing shared and distinct processes helps elucidate how mutations like *Kras* alter normal cell-state transitions and initiate oncogenic programs.

### Leukemic derailment in AML initiates from immature cells

Next, we applied Decipher to investigate the complex and poorly understood derailment of early leukemic cells in AML. We collected 104,116 single-cell transcriptomes from bone marrow specimens of a cohort of AML patients bearing *TET2* epigenetic mutations (*n* = 12), with and without *NPM1* mutations, as well as a healthy donor as reference (**Supplementary Information; Table 3**). *NPM1* is among the most commonly mutated genes in AML (20-30% of cases), yet its role in leukemogenesis [57] is unknown. *NPM1*^mut^ AML often co-occurs with mutations in the epigenetic modifiers *TET2* and *DNMT3A*, which are known drivers of clonal hematopoiesis, a condition associated with an elevated relative risk of progression to myeloid malignancy in older adults [58]. We know that these epigenetic mutations likely originate in pre-leukemic hematopoietic stem cells (HSCs) [59], facilitating AML development following *NPM1* mutation [60]. However, the transcriptomic consequences of *NPM1* mutation and the influence of pre-existing epigenetic abnormalities remain unclear.

Consistent with prior studies [20,61], we found significant inter-patient heterogeneity in leukemic blast cells (**Fig. S2a,b**), which is unlikely due to technical effects given that lymphocytes are well mixed. Surprisingly, we found that the most immature, HSC-like cluster (448 cells; 0.4% of total) is the top non-lymphoid cluster shared by most patients (**Fig. S2a,c**). The phenotypic similarity of immature leukemic or pre-leukemic cells across patients contrasts with the striking heterogeneity of leukemic cells, motivating the reconstruction of patient-specific trajectories diverging from normal HSPCs (**Fig. 4a**). However, our samples contained too few HSC-like cells to effectively characterize *NPM1*-mediated derailment; thus, we identified two differentially expressed surface markers from our cohort data, CD34 (log fold change: 6.67, adjusted *p* < 1e-6) and a novel maker, PROM1 (CD133[62]; log fold change = 7.23, adjusted *p* < 1e-6), and used them to enrich the immature population in *NPM1*^mut^ patients. Sorting for cells expressing either marker expanded the target population from 179 to 13,210 cells in patient AML1 (**Figs. 4b**), for a total of 29,266 immature cells from AML1–AML3 (Methods). *NPM1* mutations can be detected directly from scRNA-seq data[63] because the vast majority occur at the 3’ end of the gene[57] in AML (**Supplementary Table 1**). The expanded HSC-like population revealed cryptic heterogeneity in both *NPM1* mutation status and maturation (**Fig. S2d,e**). Our data spanning leukemic progression, especially the rare early stages around *NPM1*^mut^-mediated derailment, thus poised us to ask exactly when and how cells diverge from myeloid differentiation in normal HSPCs.

**Figure 4.**
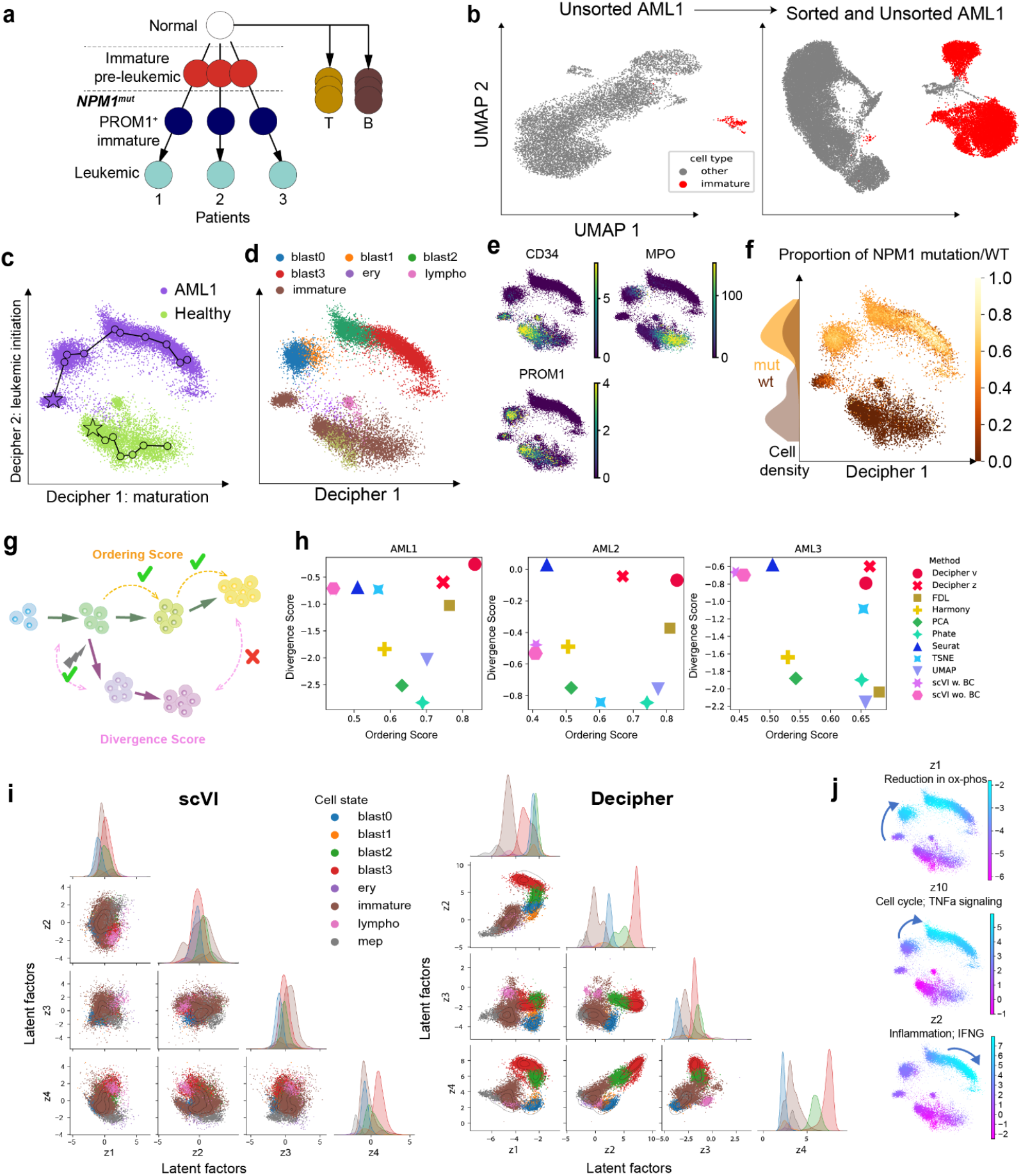
Decipher reconstructs derailed myeloid developmental trajectories in AML. **(a)** AML is characterized by patient-specific trajectories with similar immature wild-type cells, shared initial states, and divergent trajectories to terminal states following *NPM1* mutation. **(b)** UMAP projection of AML1 single-cell transcriptomes with (right) and without (left) the inclusion of sorted CD34^+^/PROM1^+^ cells to enrich for HSC-like cells. **(c–f)** Decipher space embedding of 37,395 sorted cells from patient AML1 and a healthy donor, colored by the sample of origin (c), cell states (d), key cell markers (e), and *NPM1*^mut^-to-*NPM1*^wt^ proportion (f) (Methods). Decipher 1 organizes cells along maturation and Decipher 2 along leukemia initiation axes. Lines and circles represent post-analysis trajectories; stars indicate initiating states. **(g,h)** Two metrics that measure a latent space’s interpretability and faithfulness to underlying biology (g; Methods), used to benchmark Decipher against dimensionality reduction and harmonization methods (h). **(i)** Comparison of scVI[37] (left) and Decipher (right) latent spaces for pairs of the first four latent dimensions (the other latent dimension pairs give similar results). The scVI latent space collapses biological differences while Decipher preserves them. **(j)** Decipher space colored by latent factors z1, z10, and z2, each capturing a different state transition.

### Decipher reconstructs maturation and derailment in AML

We applied Decipher to integrate data from a healthy individual with data from each patient AML1, AML2 or AML3 separately (**Fig. 4c, S3)**. For each patient, we found that Decipher 1 and 2 faithfully represent the shared processes of cell maturation and disease derailment, respectively. Specifically, Decipher 1 captures the stepwise maturation of leukemic cell states from immature to blast 0 through 3 in a leukemic derailment trajectory (**Fig. 4d, S3**), and loss of stemness and myeloid differentiation, as determined by loss of *CD34* and gain of *MPO*, in a normal cell-state progression trajectory (**Fig. 4e**). Decipher 2, in contrast, represents an axis of leukemic initiation and progression that can be further interpreted using *NPM1* mutation status. We found a subset of pre-leukemic immature *NPM1*^wt^ cells close to healthy HSCs, and an *NPM1*^mut^ progenitor-like population (blast0,1) that lies between *NPM1*^wt^ and leukemic cell states resembling myeloid-committed cells (blast2,3) in all three patients (**Figs. 4d,f** and **S3**). The increase in *NPM1*^mut^ cell fraction and upregulation of *PROM1* along Decipher 2 confirm that it distinguishes leukemic from normal cell states (**Fig. 4e,f**). Thus, similar to the pancreatitis example, the Decipher 2D space represents major axes of biological variation and preserves global relationships between cell states in the complex context of AML, correctly placing *NPM1*^wt^ leukemic cells closest to normal (**Fig. 4f**) and ordering leukemic blasts by maturation (**Supplementary Information**).

Indeed, Decipher identifies trajectories with both shared and distinct features for patients AML1 to AML3, whereas AML2 has a larger gap between *NPM1*^wt^ and *NPM1*^mut^ cells in Decipher space (reinforced by the absence of detected blast0 cells in this patient), and branching in AML3 occurs during blast1 rather than before blast0, suggesting later derailment in this patient than in AML1 or AML2 (**Fig. S3**). Such differences argue for developing personalized AML models based on patient-specific disease trajectories.

To further characterize early disease processes, we used gene set enrichment analysis (GSEA) on cells projected onto the Decipher 2 derailment axis and identified TNFa signaling, inflammatory response, IL6/JAK/STAT3 signaling, IFNg response, and KRAS signaling pathways (**Table 4** and Methods). These findings agree with the well-elucidated role of *Tet2* in repressing IL6 transcription [69], and with the association of *Tet2* loss-of-function with the accumulation of inflammatory myeloid cells in conditions from clonal hematopoiesis-related atherosclerosis [70] to AML [71].

In contrast to Decipher, we found that the visualization approaches tSNE [65], UMAP [40,48] and force-directed layout (FDL) [40] fail to capture the global data geometry, the expected overlap in immature cell states, or the order of blast maturation stages. On the other hand, data integration methods tend to force cell states—including leukemia cells and normal HSPCs—to overlap, and thus cannot be used to characterize derailed differentiation (**Fig. S4**). To systematically benchmark the ability to characterize derailed trajectories, we defined two metrics that evaluate biological faithfulness to AML derailment (**Fig. 4g** and Methods). Ordering score evaluates whether each method’s latent space correctly orders cell states by maturation stage. Divergence score assesses the preservation of biological differences by rewarding immature cell proximity and penalizing terminal state mixing between normal and disease samples. We applied these metrics to a range of visualization (FDL[40], UMAP[40,48], tSNE[65], PHATE[49]), dimensionality reduction (PCA), and integration methods (Seurat[31], scVI[37], Harmony[66]). Decipher components and latent factors scored highest in both metrics for all three patients, demonstrating that it balances between integrating across conditions and preserving their unique geometries and cell states (**Fig. 4h**).

### Decipher can represent correlated biological mechanisms

The hierarchical dimensionality reduction in Decipher confers more expressive factors that cannot be attained by simply increasing the number of latent factors in other methods [45]. Since Decipher’s latent factors can be correlated (**Fig. 4i**), they are able to capture overlapping transcriptional programs between trajectories and shared mechanisms underlying consecutive cell state transitions. For example, z7 is mainly encoded with Decipher 1 (**Fig. S5a**) and represents common trends along both normal and AML trajectories (**Fig. S5b**).

Latent factors can also highlight features that distinguish normal and perturbed states. For example, the first two factors (z1, z2) are correlated in immature cells but not in blasts (**Fig. 4i**), and represent different transitions along AML derailment**—**z1 is highest early in blast formation (blast0, 1 and 2), z10 marks an intermediate (from blast1 to 2), and z2 marks the final stage of leukemic maturation (blast3) (**Fig. 4j**). GSEA reveals that z1 is enriched for reduction in oxidative phosphorylation, a pathway that is altered in leukemic stem cells[67], while z10 and z2 are enriched for TNFɑ, IFNg and inflammation[68,69] (**Fig. 4j** and **Table 4**). Indeed, the enrichment of IFNg and inflammation in z2 (**Table 4**) is expected as it captures most mature myeloid/monoblastic cells [68,69]. Decipher thus enables a more comprehensive and nuanced understanding of cellular dynamics, especially when integrating normal and perturbed conditions.

Decipher’s unique ability to model correlated latent factors avoids the requirement for independent factors, which can remove biological differences between cell states. For example, scVI collapses the healthy and AML conditions onto each other in every latent dimension (**Fig. 4i**), deforming the geometry and disrupting continuous trajectories (**Fig. S4b**).

### Gene patterns along Decipher trajectories reveal altered regulation in AML

To uncover gene expression dynamics along cell maturation and disease derailment trajectories visualized by Decipher (**Fig. 4c,d**), its decoders can directly transform any cell state in Decipher space, including sparsely sampled states, to their corresponding gene expression mean and variance.

We constructed paths along trajectories in Decipher space and computed expression along these paths to obtain gene patterns (**Figs. 5a** and Methods). Trajectories can be defined manually or obtained with any trajectory inference method. We chose to implement a simple method that clusters cells in the latent space, generates a minimum spanning tree to link clusters, and then interpolates between clusters in Decipher space (**Fig. S6** and Methods). The resulting coordinates along those paths define a pseudotime called *Decipher time* (**Fig. 5a**). Importantly, since different conditions are integrated in the same joint space, the trajectories have comparable Decipher time. The gene patterns inferred by the Decipher decoder can thus be directly compared ‘out of the box’, without the additional challenging trajectory alignment required by standard integration methods. The similar patterns of key developmental markers *CD34[70], AVP* (stemness), *PABPC1* (protein synthesis in HSC differentiation) and *LYZ* (myeloid differentiation) in aligned segments of the trajectories confirm that the inferred pseudotime is comparable between datasets (**Fig. 5b**).

**Figure 5.**
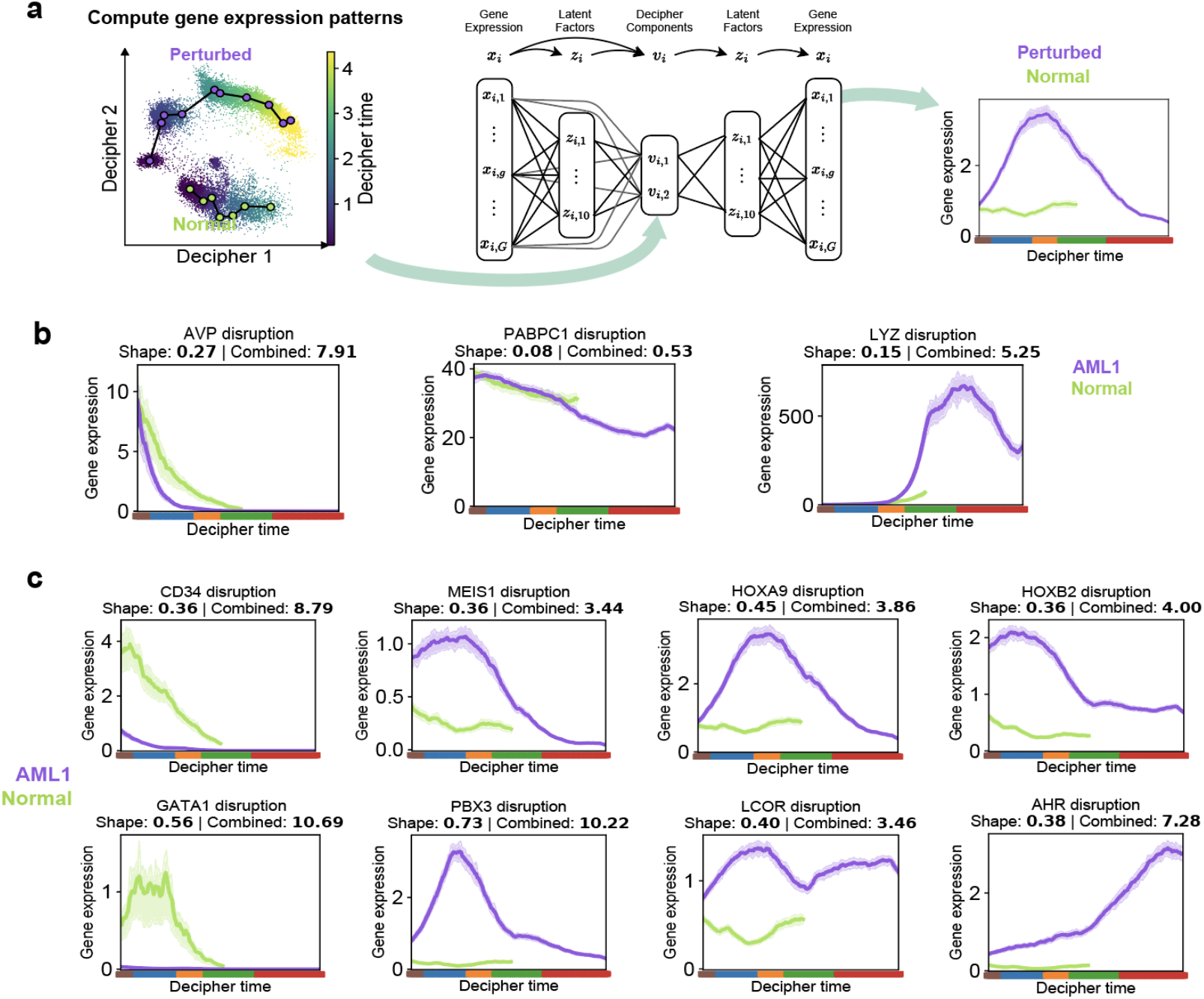
Reconstructing gene expression patterns and characterizing the regulatory landscape in AML compared to healthy HSPCs. **(a)** The Decipher generative model reconstructs gene expression along each trajectory directly from the 2D Decipher representation. **(b)** Reconstructed expression of stemness and differentiation markers for each trajectory along Decipher time. Shaded bands represent the interquartile range of Decipher model uncertainty (Methods). **(c)** Reconstructed gene expression dynamics of *HOXA9* and *MEIS1* (known deregulated TFs) and other disrupted TFs. Solid lines show inferred mean, shaded areas reflect +/-1 s.d.

To shed light on the regulatory mechanisms underlying disease derailment, we used Decipher time to estimate when transcription factors (TFs) are maximally expressed in each trajectory. By computing this for all TFs, we found that TFs are upregulated in concert at specific locations along normal hematopoiesis (**Fig. S7a**). In contrast, all AML patients display a global loss of TF coordination, including the peak at blast0, which is lost in AML precisely when *NPM1* mutations appear (**Fig. S7a,b**). At the level of individual TFs, we confirmed known upregulation of homeobox genes (*HOXA9[71], HOXB2*) and their cofactors *MEIS1[72]* and *PBX3[72]*, and downregulation of *GATA1[73]* when *NPM1* mutations appear (**Fig. 5c**). However, we also found diverse TF expression dynamics (e.g. *LCOR, AHR*), illustrating that summarizing by peak expression is insufficient to capture the complexity of transcriptional regulation (**Fig. 5c**). We thus developed a more systematic approach to quantify altered expression between two trajectories next.

### Basis decomposition reveals specific gene dysregulation in AML

To quantify the differences between gene trends along two trajectories, we devised a probabilistic framework that assumes the expression of each gene can be approximated by a weighted combination of a few representative patterns with distinct temporal dynamics, such as ascending, descending, or peaking in intermediate states (Fig. 6a and Methods). The model further assumes that representative patterns are shared between normal and perturbed trajectories, but with a different scale parameter and weights. The representative patterns are mathematically defined as *basis functions* that are simultaneously inferred with the decomposition parameters and capture dominant dynamics along trajectories. The decomposition weights (β) indicate which patterns are associated with each gene, and the scale parameter (*s*) indicates the magnitude of expression (high or low) of the pattern. Specifically, we modeled the patterns using Gaussian Processes adapted from ref.[74] and approximated them using neural networks (Methods).

**Figure 6.**
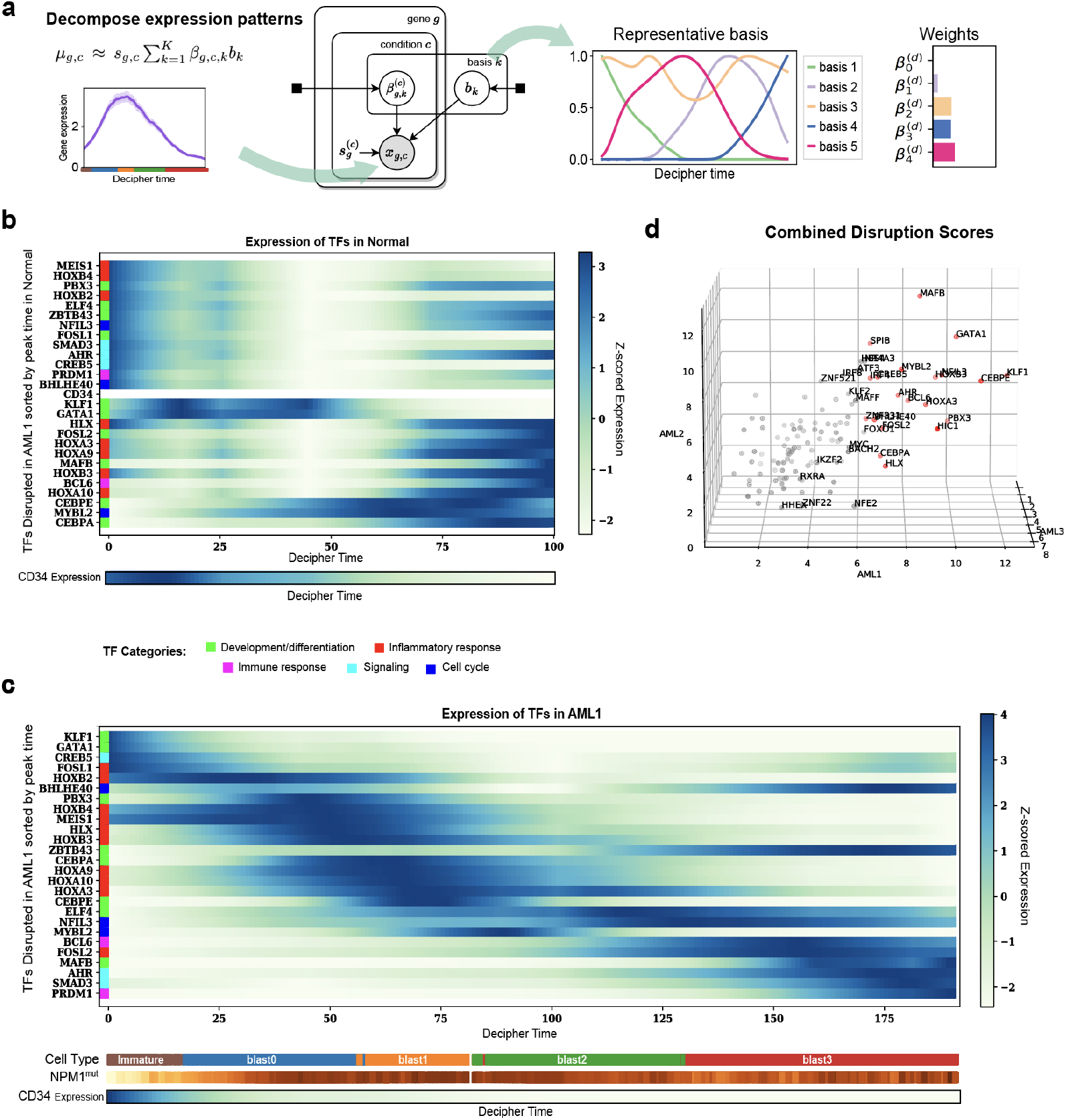
Decipher components unlock transcription factor dynamics. **(a)** Overview of Decipher’s probabilistic basis decomposition and disruption quantification method. The neural basis decomposition learns the dominant representative patterns and decomposes each gene expression pattern onto them; the coefficients on the basis for each gene are compared between the normal branch and the AML branch to compute the disruption score (Methods). **(b,c)** Timing of TF expression in normal (b) and AML1 (c) samples. Heatmaps show log-transformed and z-scored expressions for the top 20 TFs with the highest combined disruption scores in AML1, combined with known TFs from the literature, sorted by timing of maximum expression in AML1. The HSPC marker *CD34[78][72]* is included for calibration. Colorbars correspond to cell type and proportion of *NPM1*-mutated cells among the 30 nearest neighbors of each cell in AML1; both are smoothed over the 50 nearest neighbors in the Decipher space. **(d)** The combined disruption score (Methods) of TFs in three AML patients. Red points indicate TFs that are among the top 20 disrupted TFs in at least one patient.

Using the basis decomposition, we can distinguish changes in the temporal dynamics of genes from changes in the overall scale of expression. The *shape disruption* measures the distance between decomposition weights independent of the scale, while *combined disruption* considers both weights and scale (Methods). We computed shape and combined disruption scores for all genes and identified conserved (unchanged between normal and perturbed) and disrupted (altered) genes in both measures (**Fig. S8a**). For instance, homeobox genes *HHEX* and *HOXB2* have a high shape disruption score (**Fig. S8b**), while inflammatory genes *CXCL3* and *CXCL8*, as well as *KMT2A* (which interacts with Menin, a target of *NPM1*-mutated AML clinical trials[75]) have a high combined disruption score, as they are upregulated in immature AML cells compared to normal, with substantial differences in scale (**Fig. S8a,c**).

Many of the genes with highest shape disruption (e.g., *HHEX, HOXB2*, **Fig S8b**) arise early in AML, around the initiation of *NPM1* mutation, and subsequently drop to similar expression levels in advanced blasts, highlighting the importance of enriching our data with early leukemic progenitors to detect such initial events. Others, such as TNFɑ pathway (*CCL4, PHLDA1*) and oxidative phosphorylation (*ALAS1, TCIRG1*) genes, peak late and offer insight into the final transitions to disease (**Fig. S8d,e**).

The ability of disruption scores to associate genes with cell-state transitions offers an opportunity to understand regulatory events underlying derailment in AML. We collated a list of TFs with high disruption scores and known TFs from the literature, and found that they are transcribed in coordinated waves in normal hematopoiesis and as a cascade in AML (**Fig. 6b,c**). The key myelopoiesis regulators *GATA1* and *KLF1* are at the top of this cascade, followed by *HOXA9* and *MEIS1* (known to be altered specifically in *NPM1*-mutated AML[72]) at the time *NPM1* appears (**Fig. 6c**). This suggests that *TET2* mutation disrupts the function of key hematopoiesis TFs, which propagate to additional TFs in the gene regulatory network. Interestingly, the upregulation of HOX TFs upon *NPM1* mutation coincides with an increase in Interferon Type 1 signature genes[68][76] including *LY6E, FAM46C, ADAR*, and *TMEM238* (**Fig. S9a**). This is in contrast to components of Type II Interferon (IFNg) response, including MHC-II genes that are upregulated most in early immature cells (**Fig. S9b**), suggesting their link to *TET2* mutation. Moreover, we observe that genes in the proinflammatory TNFɑ pathway (**Fig. S8c**), the inflammatory cytokine gene *IL-1*, and AP-1 component *FOS* (**Fig. S9c**) are upregulated towards the end of the trajectory in transformation to blasts.

To determine whether dysregulated TF patterns can generalize, we compared the top disrupted TFs of all 3 AML patients (**Fig. S10**), and observed strong similarity in the combined disruption score across patients, with key TF genes including *HOXB3, HOXA3* and *GATA1* among the top 20 disrupted TFs in AML1 and AML2 (**Figs. 6c,d** and **S10a**). AML3 presents other disrupted regulators, such as *MYC*, which is known to be overexpressed in AML[77] (**Fig. S10b**). In all 3 patients, we observe a cascading effect of key disrupted TFs over time (**Fig. S10a,b**) that is supported by single-cell ATAC-seq data (**Supplementary Information**). Our approach thus resolves the timing of TF activity with respect to significant events such as genetic mutations and signaling pathway activation, guiding further studies of regulatory relationships.

To evaluate our disruption scores in the context of a larger cohort, we performed differential gene expression analysis between *NPM1*^mut^ AML and normal samples using a publicly available cohort of 125 *NPM1*^mut^ AML samples and 16 HSC-enriched normal subpopulations (see **Data Availability**; Methods). Examining the top 20 disrupted TFs in AML1 (combined disruption score), we found that 18 out of 20 are differentially expressed (absolute log_2_ fold-change > 1.16; *p* < 6.39e-03) in the larger cohort. We also find 12 out of the top 20 disrupted TFs in AML2 (absolute log_2_ fold-change > 1.6; *p* < 4.06e-03) and 7 out of the top 20 TFs in AML3 (absolute log_2_ fold-change > 1.52; *p* < 6.01e-09) to be differentially expressed in bulk data. Interestingly, TFs that are not detected in the larger cohort (e.g., *POU2AF1, MAFF*; **Fig. S8f**) exhibit altered expression in early immature cells whose signal would be diluted in bulk data dominated by blasts. It is noteworthy, however, that TFs disrupted in AML1 that overlap with the bulk differentially expressed are not the most enriched genes, but rather among the top 26% of genes ranked according to absolute log_2_ fold-change. For example, six *HOX* family genes (**Fig. 6b,c**) have an average rank of 14,376 out of 48,850 genes in the bulk analysis. The most differentially expressed genes reflect enrichment in terminal blasts, whereas our profiling of early immature cells and computational modeling of their dynamic gene expression deciphers regulators of leukemic initiation.

### Comparative analysis of early-occurring epigenetic mutations in AML

We applied Decipher to compare derailment mechanisms between *TET2*^mut^ and *DNMT3A*-mutated patients. *TET2* and *DNMT3A* are epigenetic regulators with opposing roles—*DNMT3A* adds and *TET2* removes methyl groups[79]. Mutations in these genes, along with *ASXL1*, are observed in pre-leukemic lesions and clonal hematopoiesis, supporting a stepwise mechanism for AML progression by which normal HSPCs acquire mutations in epigenetic modifiers prior to a transformative event such as an *NPM1* driver mutation[58]. We compared derailment in these epigenetic contexts to investigate how disease-priming mutations with opposing roles lead to similar vulnerabilities, and to determine whether they share mechanisms of leukemogenesis.

We thus profiled unsorted and CD34-sorted bone marrow cells from *DNMT3A*^mut^ patients (AML13–17), three of whom also harbor *NPM1* mutations (**Table 3**), and used our original *TET2*^mut^ cohort to annotate the maturation stages of clusters in this single-cell data (Methods). In the *DNMT3A*^mut^ patients, Decipher also successfully aligns AML maturation and normal HSPC differentiation along the Decipher 1 axis, and resolves disease derailment along Decipher 2 (**Fig. S11a–d**). We found that in the two *DNMT3A*^mut^ *NPM1*^mut^ patients with sufficient sorted cells (AML14, AML15), *PROM1* marks blast0 states (**Fig. S11b,d**), and 8 of the top 20 disrupted TFs overlap, including regulators of myeloid lineage commitment and differentiation (*CEBPE, HOXB3, AHR, KLF2, MYBL2)*, inflammatory response (*JUND*), homeobox cofactor (*MEIS1*) and *ZBTB20*. Interestingly, the reduction in oxidative phosphorylation pathway is enriched along Decipher 2 in both *DNMT3A*^mut^ and *TET2*^mut^ patients (**Table 4**).

For a more comprehensive comparison of epigenetic mutations, we identified disrupted TFs according to both shape and combined disruption (**Fig. S11e**). Many combined disrupted genes partially overlap; for example, *CEBPE* and *HOXB3* are disrupted in both *TET2*^mut^ and *DNMT3A*^mut^ patients, while interferon regulator *IRF8* shows higher combined disruption in *TET2*^mut^ and inflammatory response regulator *JUND* shows higher shape disruption in *DNMT3A*^mut[80,81]^. Decipher thus provides a framework for the unbiased characterization of patient-specific disease trajectories and for the comparative analysis of disease mechanisms between patients and genetic backgrounds.

### Decipher characterizes disease onset in gastric cancer cohorts

In addition to pairwise comparisons, Decipher can be applied to study early and stepwise transitions in disease cohorts. To illustrate this, we used scRNA-seq data from intestinal (IGC) and diffuse (DGC) primary gastric tumors and paired adjacent non-malignant tissue from 24 patients[82] (**Fig. 7a**). Each type of gastric cancer was previously shown to capture a partial disease trajectory—specifically, cell-state transitions I1 to I3 in IGC, and D1 to D3 in DGC[82]. However, visualizing the data of all patients with UMAP suggests alternate cell-state transitions that are incorrect; for example, that enteroendocrine (non-malignant) cells transition to I3/D3 states before I2/D2 and I1/D1 (**Fig. 7a** and ref.[82]).

**Figure 7.**
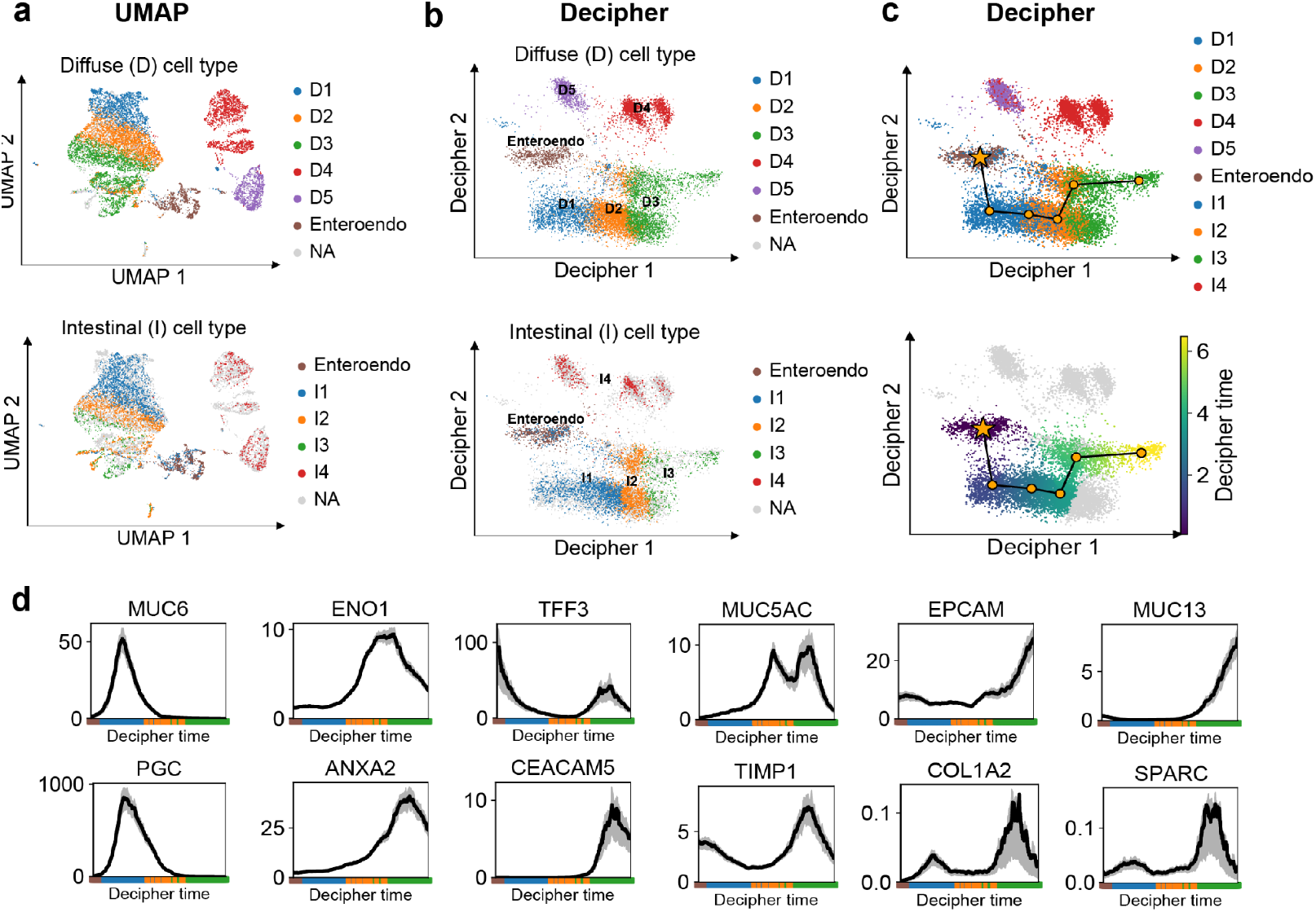
Decipher aligns gastric cancer onset. **(a)** 2D UMAP projection and trajectory inference with Monocle applied to scRNA-seq data from 24 gastric cancer (GC) and precancerous lesions[82]. Single-cell data from both intestinal (IGC) and diffuse (DGC) types were merged to infer a joint tree using Monocle3[83] (in gray line). The order of disease progression states (I1-I3 in IGC and D1-D3 in DGC) is not captured by the tree. **(b-c)** Decipher reveals the order of cancer stages, while still harmonizing the different types of cancer without requiring a batch correction method or a dimensionality reduction method for visualization. We separately plot D (top) and I (bottom) cells, but Decipher is trained on all cells. Cells in Decipher space are colored by cancer progression stages in DGC and IGC (b), and inferred Decipher trajectory and Decipher time on merged data is shown in (c). **(e)** Decipher’s reconstructed gene patterns for known markers of progression states along the shared trajectory.

We pooled all patients together and used Decipher to derive a more straightforward and interpretable representation of tumor progression in the two GC types. The Decipher 1 axis correctly aligns cells along a continuous shared progression, with non-malignant enteroendocrine, I1 and D1 cells at the start, followed by I2 and D2 intermediate states, and finally I3 and D3 malignant states (**Fig. 7b**). The inferred Decipher trajectory (**Fig. 7c**) reveals the upregulation of malignancy-associated genes at the correct cell-states, including *MUC6* and *PGC* in normal gland mucous cells, *ENO1* in intermediate state I2 and *MUC13* and *CEACAM5* in malignant states (**Fig. 7d**). The alignment of GC types also illuminates the relative timing of disease progression, predicting that premalignant cells transform to malignancy in IGC later than in DGC, as I1 extends further than D1 along Decipher 1 and Decipher time. The Decipher 2 axis, on the other hand, separates the diffuse-like malignant state (D4) in DGC, illustrating the drastic derailment from D3 upon upregulation of key DGC markers such as *COL1A2* in D4. Finally, fibroblasts and endothelial cell states (I4, D5), which are not a part of the progression of cancer, are preserved as distinct states, highlighting Decipher’s ability to represent both continuums and distinct states.

## DISCUSSION

Decipher is a deep generative model designed to learn and visualize joint representations of normal and perturbed data. Whereas single-cell data analysis approaches carry out latent factorization and 2D projection as distinct steps, Decipher is unique in merging the two within a single probabilistic, hierarchical structure. As a result, Decipher not only provides direct 2D visualization, but also captures more intricate information while remaining interpretable and discovering dependencies between the underlying latent factors. In addition to visualizing cell states with less distortion than other methods, Decipher can use the joint representation to infer trajectories of cell-state transitions and identify genes with disrupted expression patterns using a novel basis decomposition technique. Decipher scales to large cell numbers due to its VAE model formulation, which allows for stochastic variational inference[84]. Other approaches deploy VAEs [37,42] or Gaussian process latent variable models (GP-LVMs) [85,86] for nonlinear dimensionality reduction, but neither are capable of simultaneous 2D visualization, and GP-LVMs do not scale to large datasets.

In simulated data, Decipher more accurately preserves sparsely sampled cell-state trajectories and better maintains the geometry of the data than other methods. We anticipate that Decipher will be a valuable tool for discovering how perturbation or disease initiation derail development. It successfully separates normal and mutant cell trajectories in a mouse model of PDAC bearing a mutation in the tumorigenic driver *Kras*, revealing the activation of distinct molecular pathways in response to oncogenic stress. Decipher’s broad applicability is also evinced by its successful joint mapping of transitions from premalignant to malignant cell states in two subtypes of gastric cancer.

The early stages of tumor initiation are understudied in primary AML, and findings from animal models only partially translate to humans [87]. AML presents significant genomic and transcriptomic heterogeneity, suggesting multiple vulnerable states and origins of derailment from normal hematopoiesis [20][88]. Decipher is able to characterize patient-specific divergence from normal myeloid differentiation, confirmed by *NPM1* genotyping, whereas other integration methods distort the global geometry of trajectories. Our work discovered and characterized a rare subset of *PROM1*+ cells in *NPM1*-mutated samples that likely define a pre-leukemic cell population [89][90]. Decipher also revealed that *NPM1* mutations trigger the upregulation of inflammatory genes and IFN responses, following loss of coordinated myeloid TF expression due to *TET2* mutations. These findings are consistent with studies linking high *HOX* expression to mutant *NPM1* and its aberrant cytoplasmic localization in leukemic persistence [91][92].

Recent studies in mice demonstrate that loss of *Tet2* induces the expansion of aberrant inflammatory monocytic populations, by establishing a pro-inflammatory microenvironment[[68][93]. Similarly, we find the upregulation of IFN type 2 (specifically, MHC-II) genes in early *TET2*-mutated cells in primary samples. *NPM1* has been reported to regulate IFNg-inducible genes in HeLa cells [94], but the link is not established in AML. Our pseudotime-resolved characterization of transcriptional dynamics shows genes involved in IFN type 1 response to be highly expressed, specifically in transition to aberrant *NPM1*-mutated progenitor cells, coinciding with the expression of *HOX* TF genes. In further transformation to blasts, we observe the upregulation of genes encoding TNFɑ, IL-1, and FOS. The diverse patterns of chemokines and cytokines along the leukemic transformation trajectory also point to possible dysregulated interactions among them [95]. Inflammatory cytokines such as IL1 can indeed regulate hematopoietic stem cells and promote disease progression in models [96]. While our data does not contain significant non-leukemic myeloid populations and cannot resolve the cellular source of the IL6 and other cytokines responsible for inducing these programs, their stark upregulation along the Decipher 2 component supports work in *Tet2* murine models [72], suggesting this inflammation drives cellular plasticity enabling leukemogenesis, rather than merely being coincident to it [71–75].

In addition to its role in shaping the AML microenvironment [93], cell-intrinsic inflammation is induced by *NPM1* perturbation in mice, leading to myelodysplastic syndrome-like phenotypes [97] and driving progression to AML[98]. These observations motivate future studies on inflammatory response as a consequence of *NPM1* perturbation, compared to epigenetic remodeling in clonal hematopoiesis, and studies disentangling the role of pre-existing epigenetic mutations in inducing an inflammatory environment crucial for disease transformation. Extending the application of Decipher to other primary cancer samples as well as animal models can guide therapeutic strategies for modulating the TME and cell-intrinsic effects by attenuating the inflammatory response and, in turn, inhibiting cancer progression or increasing sensitivity to treatments. Decipher could also be extended to characterize multimodal datasets more effectively.

## DECLARATIONS

### Ethics approval and consent to participate

The genomic sequencing study used in our Acute Myeloid Leukemia analysis was approved by the Institutional Review Board/Privacy Board-B of Memorial Sloan-Kettering Cancer Center (Protocol #17-549) on November 4, 2017.

### Consent for publication

Not applicable.

### Availability of data and materials

The data discussed in this manuscript will be deposited in the National Center for Biotechnology Information’s Gene Expression Omnibus (GEO) upon publication. The bulk RNA-seq data for AML patients is publicly accessible at GEO with accession IDs GSE106272, GSE49642, GSE52656, GSE62190, GSE66917, and GSE67039. Bulk RNA-seq for normal HSC-enriched subpopulations is accessible at GEO ID GSE48846.

Decipher is available at https://github.com/azizilab/decipher and has been deposited at DOI: 10.5281/zenodo.10079999. The specific code that produced the results and figures of this manuscript is available at https://github.com/azizilab/decipher_reproducibility and deposited at DOI:10.5281/zenodo.14042470. The Decipher model has also been implemented as part of scvi-tools: https://scvi-tools.org.

### Authors’ contributions

A.N., J.L.F., D.P., E.A. conceived the study and wrote the manuscript. A.N., J.L.F., D.B., D.P., E.A. designed and developed Decipher. V.P.L., D.P., and E.A. designed AML experiments and prepared samples. V.K., I.M., R.L.B., S.E., and R.L.L. performed and assisted with single-cell genomics data acquisition experiments. A.N., J.L.F., V.P.L., C.B., J.C., J.W., L.S., A.E.C, D.P., E.A. analyzed and interpreted data.

## Acknowledgements

We thank Guy Sauvageau from the Leucegene group and Josée Hébert from the Banque de Cellules Leucémiques du Québec (BCLQ) for providing clinical samples. The BCLQ is supported by grants from the Cancer Research Network of the Fonds de recherche du Québec–Santé (FRQS). We are thankful to Tal Nawy, Benjamin Izar, José McFaline-Figueroa, Manu Setty, Cassandra Burdziak, and Sopho Kevlishvili for helpful feedback and discussions.

## METHODS

### The Decipher framework

Given a dataset of single-cell gene expression (*x*_*i,g*_) of *N* cells and *G* genes, Decipher models the expression of genes in cells by learning multiple hidden representations of each cell, at increasingly finer detail: the Decipher components *v* gives a high-level two-dimensional representations, and the ten-dimensional Decipher latent factors *z* are more refined representations of cells. To achieve this, Decipher successively combines multiple neural networks in a probabilistic framework and successively generates hidden representations with increasing dimensions (**Fig. 1b**).

Decipher is a generative model that extends traditional Variational AutoEncoder models (such as scVI and others [24, 25]) by adding an additional neural network, encoding a higher level two-dimensional latent space on top of their latent space. Importantly, this extra layer allows Decipher to model more complex cell state distributions with potentially dependent latent space factors; it provides an interpretable decomposition of the latent factors and a ready-to-use visualization of the data. Moreover, the top two-dimensional Decipher components enable visualization of the space directly from the model without additional dimensionality reduction. To fit Decipher to data from multiple samples, the samples are simply concatenated.

Fitting the probabilistic model defined by Decipher is done with amortized variational inference [5]. In other words, the Decipher model and the Decipher inference together form a special kind of Variational Auto-Encoder [21], with two nested encoders and two decoders, allowing it to encode and decode from any level of representation to any other level of representation (**Fig. 1b**), which is a novelty of our approach.

The Python code implementing our method is available at https://github.com/azizilab/decipher. For each method presented below, we reference its corresponding Python function. Our Python code follows the architecture of the `scanpy` package [43], with computation functions in a `.tl` submodule and the plotting functions in a `.pl` submodule. In the code snippets of the methods below, we assume that we have imported the `decipher` package as follows `import decipher as dc` and that the data of interest is in an AnnData object called `adata`. For instance, training Decipher is done with `dc.tl.decipher_train(adata)` and plotting the Decipher space colored by cell type is performed with `dc.pl.decipher(adata, color=“cell_type”)`. The Decipher model is also implemented in the `scvi-tools` package [12].

### The generative model

Decipher first models each cell *i* as a two-dimensional standard normal variable *v*_*i*_, termed *Decipher components*, representing the two largest axes of cell heterogeneity, such as cell type or cancer progression stages. This representation (*v*_*i*_) directly enables two-dimensional visualization of the data. A learnable neural network *f* –a *decoder* – transforms each *v*_*i*_ into a distribution over medium-dimensional vectors, a latent space representing cell state.

Decipher samples a cell state *z*_*i*_ conditionally on *v*_*i*_ from the distribution induced by f(*v*_*i*_); the *z*_*i*_ contains richer information about cell *i* than *v*_*i*_. The *z*_*i*_s are medium-dimensional. They are substantially lower dimensional than the number of genes but higher than *v*_*i*_ and can thus capture more information regarding cell variability (we set the dimension to 10 in our experiments). We refer to the medium-dimensional variables *z*_*i*_ as *Decipher latent factors*. These are comparable to the latent variables of other VAE-based or matrix-factorization-based methods [23, 25, 31].

Then, a second neural network *h* – a second decoder – with a softmax output layer, transforms the state *z*_*i*_ into the normalized gene expression values (*µ*_*i,g*_)_1*≤g≤G*_ = *h*(*z*_*i*_) expressed in cell *i*. These normalized expressions are finally scaled by the (observed) library size of the cell, *l*_*i*_, and the observed gene counts *x*_*ig*_ are sampled from a negative Binomial distribution with mean *µ*_*i,g*_ and dispersion *θ*_*g*_ specific to each gene *g*. This generative process is represented in **Fig. 1b** and is described mathematically as follows:

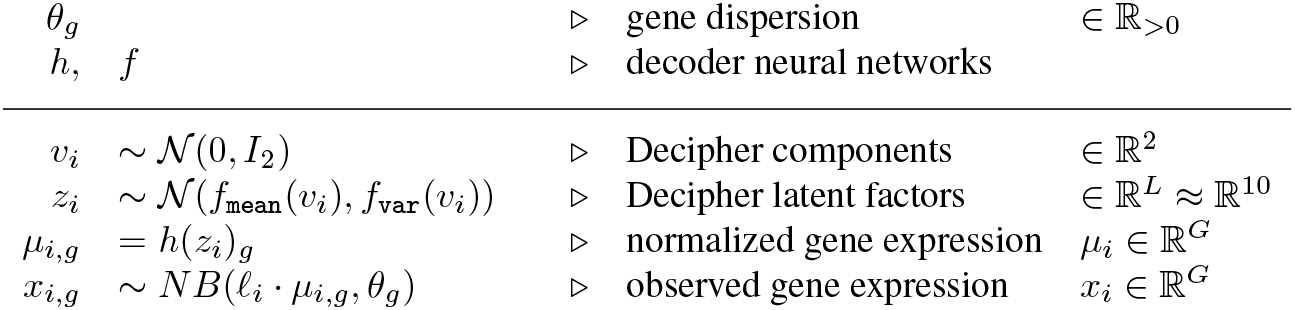

|  |  |  |
| --- | --- | --- |
| $\theta_g$ | ▷ gene dispersion | $\in \mathbb{R}_{>0}$ |
| $h, f$ | ▷ decoder neural networks | |
| $v_i \sim \mathcal{N}(0, I_2)$ | ▷ Decipher components | $\in \mathbb{R}^2$ |
| $z_i \sim \mathcal{N}(f_{\text{mean}}(v_i), f_{\text{var}}(v_i))$ | ▷ Decipher latent factors | $\in \mathbb{R}^L \approx \mathbb{R}^{10}$ |
| $\mu_{i,g} = h(z_i)_g$ | ▷ normalized gene expression | $\mu_i \in \mathbb{R}^G$ |
| $x_{i,g} \sim NB(\ell_i \cdot \mu_{i,g}, \theta_g)$ | ▷ observed gene expression | $x_i \in \mathbb{R}^G$ |

The mapping functions *f* and *h* are neural networks. In practice, Decipher uses a single linear layer for *h* as suggested by Svensson *et al*. [36]. *Meanwhile, *f* has two linear layers interleaved with ReLU activations. The last layer of *f* produces a vector in* ℝ^*2L*^ that is split to form two outputs: *f*_mean_ ∈ ℝ^*L*^ and *f*_var_ ∈ ℝ^*L*^. The choice of negative binomial distribution can be replaced by other distributions if the user believes it is more appropriate for the data at hand.

### High-level summary

The Decipher components *v*_*i*_ represent the high-level organization of the cells and form the Decipher space. This space provides a ready-to-use 2D representation of the data without requiring further projection methods such as UMAP or t-SNE. It offers direct visual access inside the probabilistic model. Then, through the neural network *f*, each *v*_*i*_ induces a cell state *z*_*i*_, a more detailed representation of cell *i*. The space of the *z*_*i*_ corresponds to the latent space of traditional variational-autoencoders.

### Inference for Decipher’s deep probabilistic model

Given the observed gene expression data 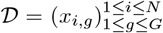 and the parameters (*θ*_*g*_)_1*≤g≤G*_ *f,h*, the Decipher’s probabilistic model defines a posterior *p* (*v, z* | *𝒟*) over the latent variables *v* = (*v*_*i*_)_1*≤i≤N*_ = (*z*_*i*_)_1≤*i*≤*N*_. We use variational inference [5, 17, 41] to approximate this exact posterior with a variational approximation *q* (*v, z*).

### The structure of the variational family

We use amortization over the local variables *v*_*i*_, *z*_*i*_ in function of the observations *x*_*i*_, such that the variational family becomes: 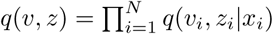 which always factorizes as

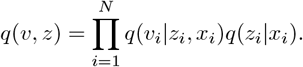

The amortized distributions are set to diagonal Gaussian distributions with parameters (mean and variance) given by neural networks – the encoders. The first neural network transforms *x*_*i*_ to the mean and the variance of the distribution *q*(*z*_*i*_|*x*_*i*_), and the second neural network transforms (*z*_*i*_, *x*_*i*_) to the mean and the variance of the distribution *q*(*v*_*i*_|*z*_*i*_, *x*_*i*_). We denote them as: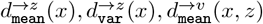, and 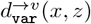, such that:

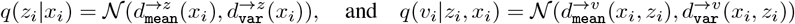

### The variational inference objective

Variational inference seeks to minimize the KL divergence between the variational posterior q and the exact posterior *p*(*·* | *𝒟*). It is equivalent to maximizing a lower bound of the evidence, called the ELBO [5]:

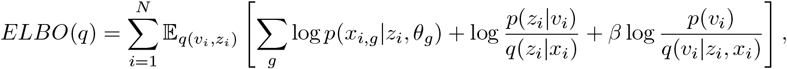

where *β* is a scalar controlling the importance of the prior *p*(*v*_*i*_), between 0 (no prior) and 1 (standard ELBO) [14].

Because we chose the variational posteriors q to be Gaussian distributions, we can reparametrize the expectations to sample unbiased low-variance estimates of the ELBO. To obtain a sample for (*z*_*i*_, *v*_*i*_) from *q*(*v*_*i*_, *z*_*i*_|*x*_*i*_) = *q*(*v*_*i*_|*z*_*i*_, *x*_*i*_)*q*(*z*_*i*_|*x*_*i*_), we first sample a reparametrized *z*_*i*_ from *q*(*z*_*i*_|*x*_*i*_) and then sample a reparametrized *v*_*i*_ from *q*(*v*_*i*_|*z*_*i*_, *x*_*i*_) [21].

The gradients are then computed using automatic differentiation. To scale up to large datasets of cells, we further subsample the outer sum using a minibatch size of 64 observations to perform stochastic variational inference [15]. We use the Adam optimization algorithm [20] to execute the gradient updates. The code is implemented in Python using Pyro [3].

Because we have little prior on the distribution of the Decipher components *v* (remember that a limitation of other methods is that the prior enforces unrealistic independence between latent variables), we set *β* to a low value 1e − 1 in our experiments.

The Decipher model can be fitted using the function `dc.tl.decipher_train(adata)`.

### Generating the Decipher space *v* and the latent space *z*

Once the inference is performed, the variational posteriors *q*(*v, z*|*x*) are fitted to the data. The “encoders” 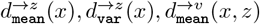 and 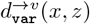 give the posterior expected values of *v*_*i*_ and *z*_*i*_ given each cell *x*_*i*_. For each cell *x*_*i*_, we compute (as in any auto-encoder architecture):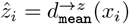 and 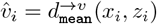.

The Decipher space *v* and latent *z* are automatically computed when calling `dc.tl.decipher_train`. They are stored in `adata.obs[“decipher_v” ]` and `adata.obs[“decipher_z” ]`.

### Rotating and aligning the Decipher space

Decipher does not use sample or batch IDs when learning the latent variables, the encoders and the decoders. However, in a post-processing step, the sample IDs (or other annotations) can be optionally used to align Decipher components to represent the most shared and most distinct information between the samples (e.g. perturbed and normal conditions), thus facilitating downstream analysis. This is accomplished by rotating or flipping the *v* components. Like most auto-encoder models (e.g., scVI [25]), the axes of the latent spaces *v* and *z* can be rotated or flipped without changing the likelihood of the data. To automatically rotate and flip the Decipher components given the user preferences, the user can specify if some cell labels should be aligned with a given component. For example, in our analysis, we choose to align the cells’ sample labels (Healthy and AML) along Decipher 2 and the cells’ cell-type labels (ordered from blast0 to blast3) along Decipher 1.

Given cell labels and their target alignment axis (e.g., ordered cell types along Decipher 1, ordered cell sample IDs along Decipher 2), we try 100 rotations (from 0 to 2*π*) and all possible axis flips (2 for *v*_1_, 2 for *v*_2_), and pick the setting that maximizes the correlations between the cell labels and their target Decipher axis.

This is accomplished by calling the `dc.tl.decipher_rotate_space` function.

### Trajectory paths construction

The Decipher components (*v*_*i*_) organize cells along visual trajectories. For instance, there are two trajectories in the joint AML-healthy data: one for healthy maturation and one for AML progression (**Fig. 4c**). The trajectories could hypothetically be traced manually by the user. Still, we propose a simple automated determination of the trajectories using, as input, the marker genes for the beginning and the end of the trajectories.

Given the cell representations (*v*_*i*_) in the Decipher space and (*z*_*i*_) in the latent space, we first cluster cells using the Leiden algorithm [37] on the latent representations (*z*_*i*_) – we use the representation (*z*_*i*_) to cluster the cells because they contain more detailed information about the cells than the (*v*_*i*_) (**Fig. S6a**, left). We then compute a minimum spanning tree between the clusters’ centroids using the distances in the Decipher space – we use the distance in Decipher space because the high-level geometry of the data is better captured by the (*v*_*i*_) (**Fig. S6a**, middle left). Finally, we use the provided marker genes to identify the trajectories’ beginning and end, from which we compute the shortest path in the minimum spanning tree (**Fig. S6a**, middle right). We use linear interpolation to form parameterized trajectories *γ* : *t ↦ v*(*t*) in the Decipher space (**Fig. S6a**, right). The time *t* that parametrizes the trajectories is called the *Decipher time*, and we compute one trajectory per sample in our analysis (*γ*_AML_ and *γ* _healthy_). If the analysis requires it, more trajectories or less could be computed.

The procedure is described in Algorithm 1 and is visually represented in Extended Data Fig. 4a. The trajectories are computed using the function `dc.tl.trajectories`.

#### Algorithm 1 Trajectory paths construction with Decipher

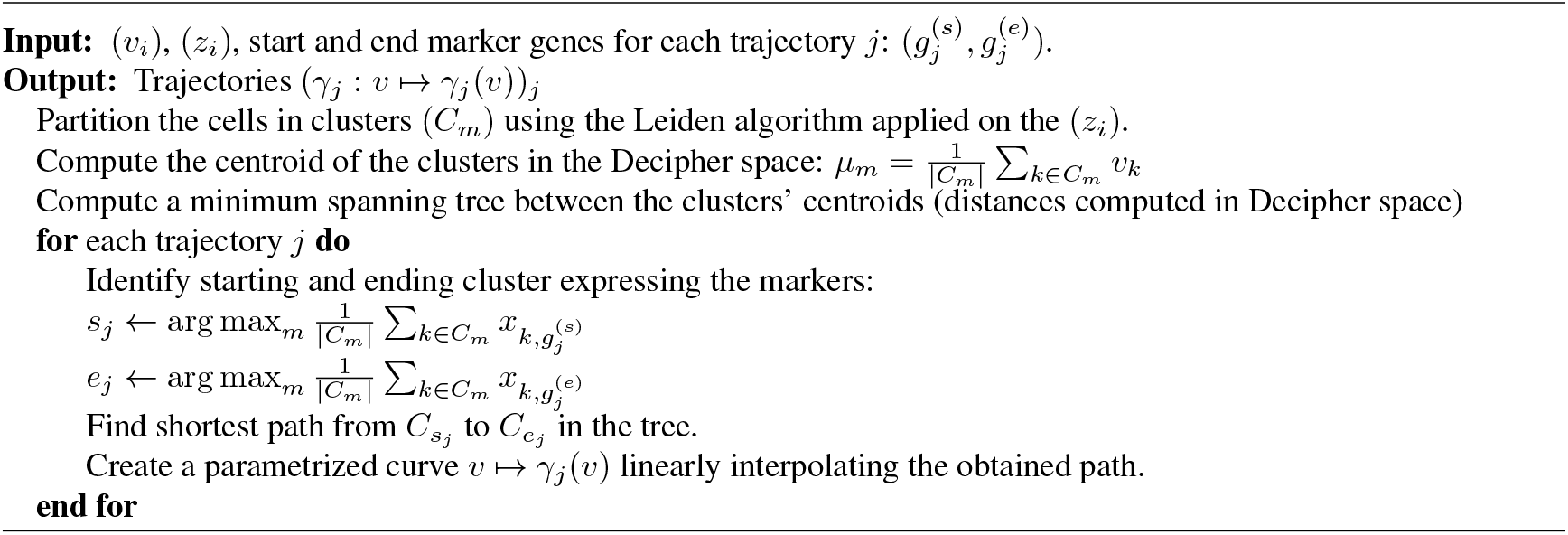

### Trajectory alignment using Decipher time

While trajectory alignment approaches e.g. with dynamic time warping [8, 18], can alter the relative lengths of trajectories by locally compressing or stretching them and potentially missing rare cell states, our jointly inferred Decipher time obtains a common time axis for both trajectories. Decipher assigns a pseudo-time value to each location along the trajectories, called the Decipher time. This is determined by calculating the curvilinear coordinate *t* along each trajectory within the Decipher space. To assign a Decipher time to any cell from our data, we project the cells onto the trajectories. That is, we compute for each cell *i* in each sample *j* (healthy or AML), the closest point *γ*_*j*_(*t*^*\**^) on the dataset-specific trajectory (*γ*_healthy_ or *γ*_AML_), and assign the time of this trajectory point *t*^*\**^ to cell *i* (**Fig. 1c**). The Decipher time is computed using the function `dc.tl.decipher_time`.

### Reconstruction of gene expression by Decipher

Given a trajectory *γ* of Decipher components, we can obtain the gene expression along this trajectory using the decoder neural networks in the Decipher probabilistic model. Indeed, we recall that the encoders and the decoders in the Decipher model can convert any gene expression into Decipher components and vice versa. Mathematically, given some Decipher components on the trajectory *v* := *γ* (*t*), we use the decoders to compute the associated expected latent factors *z* := *f*_mean_(*v*), followed by the expected normalized gene expressions *µ* := *h*(*z*), that can further be scaled up to the desired library size by multiplying it by *l*. Because Decipher is a probabilistic model, it is also possible to obtain probabilistic gene expression samples instead of a single estimate. This is particularly useful to obtain some model uncertainty around the reconstructed gene expression (**Fig. 5a**). To achieve this, we sample multiple latent factors given the Decipher components as *z*_*m*_ *∼ 𝒩* (*f*_mean_(*v*), _var_(*v*)) (instead of just *f*_mean_(*v*)) and we compute their expected gene expression *µ*_*m*_ = *h*(*z*_*m*_).

The Decipher gene expression patterns are computed using the function `dc.tl.gene_patterns`.

### Reconstruction of gene expression by scVI

Like Decipher, scVI [12] has a latent space and a decoder that can reconstruct gene expression from a latent representation z. Instead of having a low-dimensional space *v* that jointly represents the healthy and disease sample, scVI is given the label (healthy or disease) as a batch variable. Once the model is learned, scVI produces batch-corrected gene expression data for each sample by manually changing the batch label from one sample to the other, as described in https://docs.scvi-tools.org/en/stable/api/reference/scvi.model.SCVI.html#scvi.model.SCVI.get_normalized_expression (accessed on July 10, 2023).

### Basis decomposition

To quantify the difference in patterns between two trajectories, we need a metric that accounts for the temporal order of cells – two genes may have the same mean expression value but opposite patterns, e.g., ascending versus descending along trajectories. Existing methods that encode temporal dependencies are limited in modeling assumptions and scalability. For instance, tradeSeq [39] performs trajectory-based differential expression; however, approximating gene patterns with splines may not be appropriate for complex transcriptional programs, e.g., with cascades of mutations leading to cancer. Additionally, relying on built-in denoising limits compatibility with preprocessed data (e.g., from VAEs). Methods such as DPGP [27] utilize Gaussian Processes model distributions of all functions over time; however, they are computationally expensive. This motivates a metric that not only accounts for the order of cell states and uncertainty but is generalizable and scalable to the size of standard single-cell datasets. For this, we develop a method that decomposes every gene pattern on a basis of dominant patterns learned with neural networks. Comparing different patterns then becomes a comparison of their basis weights.

#### The data

The input data for the basis decomposition method is the collection of gene patterns observed over a time or pseudotime axis in different conditions, such as normal, disease, or perturbed (**Fig. 5a**). Similar to the trajectory *ϕ* : *t* ↦ *ϕ*(*t*) =: *v* that maps pseudo-time to Decipher components, we define a gene pattern *µ*_*g,c*_ for a gene *g* under condition c to be the function *t* ↦ *µ*_*g,c*_(*t*) that represents the expected expression of gene *g* at time *t* in condition *c*. With Decipher, these can be the gene expression patterns reconstructed from the inferred trajectories. In a more general setting, these patterns can be obtained with time-series bulk RNA-seq or trajectory inference methods [34, 38], applied in advance to single-cell RNA-seq data, or other dynamic features, such as chromatin accessibility (measured with ATAC-seq) or protein expression (CITE-seq).

The set of all genes is *G*, and the set of conditions is *C*. For simplicity of the exposition, we restrict the conditions to *C* = *{*healthy, disease*}*. The input data is the collection of functions *𝒟* = (*µ*_*g,c*_)_*g∈G,c∈C*_. The data has |*G*|*·* |*C*| observations, each of which is a function. In the proposed probabilistic model, each pattern *t* ↦ *µ*_*g,c*_(*t*) is considered a single observation.

#### The model

In light of commonly used generative models [4], the proposed model is a linear factor model operating in function space. For gene *g* and condition *c*, the model associates the data point *µ*_*g,c*_ with a latent scalar *s*_*g,c*_ – the *gene scale* – and a latent vector *β*_*g,c*_ of *K* dimensions – the *gene shape*. Each dimension *k* corresponds to a latent basis pattern *t ↦ b*_*k*_(*t*). The *β* _*g,c,k*_ are coefficients for the function *µ*_*g,c*_ in this basis. The coefficient *s*_*g,c*_ is the intrinsic scale of gene c in condition g, which will scale up or down the pattern computed from the bases, which, in contrast, are constrained to be between 0 and 1. The observations *µ*_*g,c*_ and the latent basis *b*_*k*_ are functions. The scale *s*_*g,c*_ and the weights *β*_*g,c,k*_ combine the latent bases *b*_*k*_ to generate the observed function *µ*_*g,c*_. That is, informally,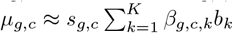. Each basis function *b*_*k*_ forms a representative pattern shared by multiple genes.

#### The weights *β*_*g,c,k*_

To ensure the interpretability of the weights, the model mimics methods like mixture or topic models [6] and draws positive weights that sum to 1. With this, a non-zero weight *β*_*g,c,k*_ signifies that gene *g* in condition c exhibits the representative pattern *k*. Specifically, the weights vectors *β*_*g,c*_ are drawn independently from a Dirichlet distribution with concentration parameter [*η, η*, …, *η*], denoted as Dir(*η*), and with density

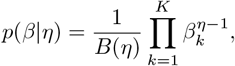

where *B*(*η*) = Γ (*η*)^*K*^/Γ (*Kη*) and Γ is the Gamma function. In terms of negative log-likelihood, this prior induces a regularization of the coefficients that will lead to sparse *β*_*k*_ when *η* < 1 and *β* closer to 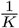 when *η* > 1. We choose *η*< 1 to associate the basis with dominant patterns and thus obtain a better interpretability of the basis.

#### The function basis *b*_*k*_

To sample the basis functions *b*_*k*_, we represent them as neural networks and sample each *b*_*k*_ by drawing its neural network parameters. Concretely, each basis function is modeled by a one-dimensional neural network with two hidden layers of 32 units each, followed by the `tanh` activation. The neural network *b*_*k*_ is of the form

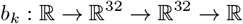

and its parameters are denoted *ϕ*_*k*_. Each *ϕ*_*k*_ is sampled from a centered diagonal normal distribution, and the variance of each of its coordinates is set to the inverse of the input dimension of the linear layer in which it appears. The study of infinite neural networks in Neal [28] demonstrates that with wide hidden units (here 32 ≫ 1) and such a prior on the parameters, the induced prior in function space is close to a Gaussian process. Gaussian processes are used in other methods [27] but are hardly scalable. Using neural networks for efficient computations solves the problem. The induced prior in function space is denoted by *φ*. Finally, to design more interpretability for the gene scale *s*_*g,c*_, we normalized the sampled basis by their maximum value so that the maximum value reached by a basis is 1.

#### The gene scales *s*_*g,c*_

The gene scales *s*_*g,c*_ are learned as variational parameters (no variational distributions), as the gene scales can greatly vary between genes.

#### The observations *µ*_*g,c*_

Finally, the gene pattern *µ*_*g,c*_ is generated from a distribution parameterized by ∑ _*k*_ *β*_*g,c,k*_*b*_*k*_ and the scale *s*_*g,c*_. More specifically, *µ*_*g,c*_ is sampled from a Gaussian process^1^ with mean *s*_*g,c*_ *·* _*k*_ *β* _*g,c,k*_ *b*_*k*_ and with a white Gaussian noise kernel 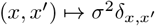 of variance *σ*^2^.

The generative process is represented graphically in **Fig. 5a** and proceeds as follows:

1. For each factor dimension *k* ∈ ⟦1, *K*⟧, draw a basis function *b*_*k*_ from the function prior: *b*_*k*_ ∼ *φ*(*b*_*k*_) (that is draw weights *ϕ*_*k*_ according to the prior detailed above)
2. For each gene *g*, and condition *c* do:
  a. For each factor dimension *k*, draw the basis weight *β*_*g,c,k*_ *∼ ℰ*(*η*)
  b. Draw the observed function *µ*_*g,c*_ from 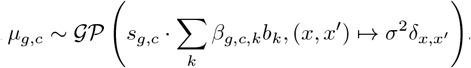.

For simplicity of notations, the *ϕ*_*k*_ are grouped in parameter *ϕ*, and the *β*_*g,c,k*_ into parameter *β*.

#### The inference

We learn an approximate posterior on the model variables *q*(*b*_*k*_, *β*_*k,g,c*_, *s*_*g,c*_) using variational inference implemented in the Python probabilistic modeling library Pyro [3], with automatic guides.

The basis decomposition is computed using the function `dc.tl.basis_decomposition`.

#### The disruption scores

From the inferred model parameters, we design multiple disruption scores that inform us of different types of disruptions for the same gene across two conditions *c*_1_ = healthy and *c*_2_ = disease.

- The scale disruption highlights the difference in gene scale between the two conditions, e.g., a gene that is up-regulated in one of the conditions. It is defined as 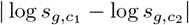.
- The shape disruption highlights the difference in gene shape between the two conditions, e.g., a gene that activated later in one of the conditions and earlier in another. It is defined as 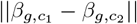.
- The combined disruption is a combination of both disruption scores to capture a general high-level disruption score, including both shape and scale. It is defined as 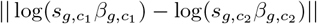.

The disruption scores are computed using the function `dc.tl.disruption_scores`.

## Data generation

### Simulated data

To evaluate Decipher’s ability to identify cell state evolution trajectories within its latent space, we simulate data with ground-truth trajectories **Fig. 2**, fit the Decipher model on this data, and evaluate the quality of the trajectory reconstruction.

We simulate data in several steps:

1. Sample random locations along the 2d *continuous* trajectories of **Fig. 2a**.
2. Remove some of the locations up to a certain percentage to simulate rare/low-sample cell state transitions: 100% in **Fig. 2a**, 90% and 95% in **Fig. b** and a varying percentage in **Fig. c**.
3. The remaining locations are denoted (*z*_*i*_) and are the ground truth cell states.
4. In particular, we consider the first coordinate of *z*_*i*_ to be the pseudotime of the cell, noted *t*_*i*_ = *z*_*i*,1_.
5. Randomly perturb the ground truth cell states to simulate noise 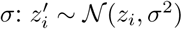.
6. Sample randomly the weights and biases of a neural network *f* with output dimension of size d (in our experiments d = 500). The neural network is used as a random function to nonlinearly transform the cell state into higher dimensional gene expression.
7. Use this neural network to map the ground truth cell states to the synthetic gene expression: 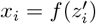.

With this simulation, we obtain random gene expression data (*x*_*i*_) that is organized along an underlying continuous trajectory with possible rare transitions.

### Evaluation metric on simulated data

We evaluate the quality of a latent space 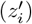 using the global preservation metric presented in Chari & Pachter [9]. We present it here briefly. Our simulation provides ground-truth cell states (the *z*_*i*_). We cluster those cell states into 20 clusters using k-means. We denote *C*_*j*_ the indices of cells in cluster *j*. Then, the global preservation metric from Chari & Pachter [9] computes the pairwise distances between each cluster in the ground-truth cell state space 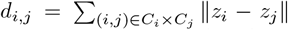, as well as in the new latent space 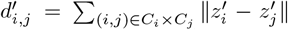. The global preservation metric is then the average Kendall-tau correlation between the distances to each cluster in ground-truth space vs new space:

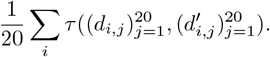

Higher is better, with a maximum of 1, indicating a perfect correlation of cluster ordering between ground truth and new latent space. The results are presented in **Figure 2c**.

### AML data collection

The TET2^*mut*^ AML cohort consists of 12 cryopreserved (DMSO) BM AML samples from the *Banque de cellules leucémiques du Québec* (BCLQ) biobank, with 10 patient specimens collected at the time of diagnosis, and two specimens from the same patient, at diagnosis and relapse (**Table 3**). The DNMT3A^*mut*^ AML cohort consists of 5 cryopreserved (DMSO) BM AML patient samples. FAB information for samples was provided by BCLQ (**Table 3**). Karyotyping, as well as mutation (variant) calling, was performed via bulk RNA-sequencing as part of the Leucegene project with data deposited on GEO with accession IDs GSE106272, GSE49642, GSE52656, GSE62190, GSE66917, and GSE67039.

All samples, as well as sorted cells, were profiled using 10X Genomics Chromium Single-cell 3’ for scRNA-seq. For scATAC-seq, cells were subjected to 10X Genomics Chromium Single Cell ATAC Reagent Kits User Guide (v1.1 Chemistry). The resulting nuclei suspension was subjected to a transposition reaction for 60 min at 37°C and then encapsulated in microfluidic droplets using a 10X Chromium instrument following the manufacturer’s instructions with a targeted nuclei recovery of approximately 5,000. Barcoded DNA material was cleaned and prepared for sequencing according to the Chromium Single Cell ATAC Reagent Kits User Guide (10X Genomics; CG000168 RevA). Purified libraries were assessed using a Bioanalyzer High-Sensitivity DNA Analysis kit (Agilent) and sequenced on an Illumina HiSeq 2500 (High Output) and NovaSeq platform at approximately 100 million reads per sample (around 5,000 nuclei) at MSKCC’s Integrated Genomics Operation Core.

#### Flow cytometry activated cell sorting (FACS)

For immature cell enrichment, FACS-purified CD34+ or PROM+ cells were subjected to single-cell RNA sequencing. Cryopreserved mononuclear cells were thawed into 10ml of prewarmed FACS buffer (phosphate-buffered saline (PBS) + 2% fetal bovine serum). Cells were pelleted at 300 × G for 5 minutes and washed again with FACS buffer. Cells were then resuspended in FACS buffer containing Human TruStain FcX(tm) (Fc Receptor Blocking Solution; Biolegend #422301) for 15 minutes at 4°C. Antibodies against CD34 (Clone 561; FITC Biolegend 343603) and CD133 (clone 7; PE Biolegend 372803) were subsequently added, and cells were stained for an additional 15 minutes at 4°C. Cells were then washed twice with 3ml of FACS buffer and resuspended in FACS buffer with DAPI. Cell sorting was performed on a Sony SH800.

## Data pre-processing and analysis

### Data pre-processing

Quantification of counts was done with SEQC [2], and 10X Genomics Cellranger [44]. Counts outputs were loaded into AnnData format using `scanpy` 1.7.2 [43]. Cells with low library size were filtered out, with the filtering threshold being selected by the knee-point of a histogram of the log10 of the total counts per cell. We obtained a median of 10,504 cells per sample and a median of 5165 molecules per cell after filtering. Data were then normalized by median library size using `sc.pp.normalize_per_cell`. Doublet detection was performed using DoubletDetection [13], with 25 iterations. scATAC FASTQ files for each sample were preprocessed to a cell-by-peak count matrix through the CellRanger ATAC pipeline [33] with modifications as described in [1].

### Annotation of AML TET2 cohort

All cells from unsorted AML samples were considered **(Fig. S2a,b)**. PhenoGraph clustering was run using 100 principal components and 15 nearest neighbors. Annotation of clusters with low counts or high mitochondrial reads was performed by visual analysis of boxplots for the log10 of counts per cluster, as well as the fraction of reads belonging to mitochondrial genes compared to all genes. We further annotated lymphoid and erythroid clusters using `scanpy`’s dotplot function to visualize key gene markers. We were able to identify these clusters by analyzing the fraction of cells in each cluster expressing key marker genes as well as the mean expression. Cells forming distinct low-count clusters and additional clusters with high mitochondrial fraction, using Phenograph clustering on a per-sample basis, were additionally removed, resulting in a global cohort of 104,116 cells.

To annotate the maturation stages of leukemic blasts, we computed correlations (`scipy.stats.pearsonr`) between the mean expression of each cluster and bulk gene expression data from sorted HSPCs [29]. The correlation calculation was limited to the 5277 most varying genes, 3475 of which overlapped with bulk data. Non-significant values (p > 0.0005) were removed (**Fig. S2c**). To control for cluster size, Shannon Diversity (**Fig. S2c**) was computed for the distribution of patient IDs in subsamples of N = 1000 (approximating the median cluster size) cells from each cluster and averaged across 20 iterations. Paired diagnosis-relapse samples (AML9, AML10) (**Fig. S2c**) annotations were considered together as they are phenotypically very similar (**Fig. S2a**). Clusters are ordered within cell-type by decreasing diversity (**Fig. S2c**).

### Mutation identification and metrics

We implemented a mutation calling protocol in order to identify mutations in NPM1 and DNMT3A. We first sorted and indexed the patient bam files using `samtools` [10], then reduced the file to the region containing the gene of interest. The NPM1 gene was analyzed from chromosome position 5:171387116-171411137, and the DNMT3a gene was analyzed from chromosome position 2:25230961-25344590. Files were then merged and indexed, then converted to FASTA format. The indexed bam file was then loaded into the Integrative Genomics Viewer [32], and the alignments were visually analyzed for the presence of the mutation. If a mutation is present, a range of 5-20 base pairs are selected for subsequent single-cell analysis (**Supplementary Table 1**). For single-cell annotation of mutations, we used the previously generated FASTA file and the mutated sequence identified for each patient to search for the presence of the mutated sequence in individual cells.

Because the mutation state may be heterozygous (a cell may have both mutant and wild-type labels), most of our subsequent analysis utilizes our defined *mutation proportion* for each cell. Since our detection of mutation is dependent on expression, which is affected by dropouts in scRNA-seq, we compute an average mutation proportion in the neighborhood of each cell. To compute the mutation proportion, we find the 30 nearest neighbors of each cell on the truncated SVD decomposition (100 components) of the normalized data. The mutation proportion for each cell is then *m*/(*m* + *w* + 1*e*^*−*10^), where m = the number of cells bearing the mutated copy of the gene and w = the number of cells bearing the wild-type copy of the gene among the 30 neighbors. The heterozygous NPM1 mutation is detected in 5-39 % of cells in each of the unsorted samples.

### Verification of immature cell enrichment in sorted samples

The primary purpose of the cell sorting was to enrich the populations of CD34+ and PROM1+ immature cells in the data (**Fig. S2d)**. To verify that this enrichment was achieved, we first performed a visual analysis of the UMAP computed on the subset of cells in AML1 originating from the unsorted collection process first, and compared it with the UMAP of cells once the sorted cells were included with the unsorted cells. We verified that enrichment of PROM1 and CD34, along with cells with low NPM1 mutation proportion, was achieved in the UMAP (**S2e**). We also quantified the expression of CD34 and PROM1 in each of four categories: immature and non-immature cells in the unsorted cells only and immature and non-immature cells in both sorted and unsorted cells. All visualization was performed using `scanpy` [43].

### Cell type mapping onto the DNMT3A cohort

To extend the annotations from the TET2 cohort to patients in the DNMT3a cohort, we combined the data for all patients in the TET2 cohort, normalizing by median library size and log transforming across the entire cohort. Cells were then grouped based on their prior cell type annotations, and 700 differentially expressed genes were identified per cell type using the T-test version of `scanpy`’s [43] *rank_genes_groups()* function. Mitochondrial and Ribosomal genes were excluded from the gene sets. Cells in the DNMT3A cohort were also combined across patients, normalized by median library size, and log-transformed. Cells were then split into clusters using PhenoGraph [22], computed using 100 principal components and k=5. We then computed the cluster centroids of the PhenoGraph clusters in the DNMT3a cohort and the cell types of the TET2 cohort by taking the mean across cells in the cluster, limiting to the differentially expressed genes. Pearson correlation (using `scipy` 1.7.0 [40]) was computed between the centroids of the two cohorts, and each DNMT3A cluster was labeled with the cell type of the TET2 cell type cluster to which it had the greatest correlation coefficient. We can then apply Decipher on these patients, following our standard analysis pipeline (**Fig. S11a**).

### Benchmarking

We evaluated the performance of Decipher on simulated data. To further benchmark the performances of Decipher on real data, we define two metrics based on our prior knowledge of AML and compare Decipher to a large spectrum of commonly used methods.

Since we do not have ground-truth trajectory values for the real data, we build metrics on prior knowledge of AML progression, AML marker genes, and our independently curated cell state annotations: immature, blast0, blast1, blast2, and blast3. Among the healthy immature cells, we further use the markers CD34 and MPO to distinguish early cells (CD34+), late cells (MPO+), and intermediary cells. Our metrics are based on the distances between annotated cell states.

- **Ordering score**: We expect the cell states in a latent space to be spatially ordered along the known cell maturation trajectories. For instance, blast1 should be between blast0 and blast2. Given the orders *o*_1_ = [immature, blast0, blast1, blast2, blast3] and *o*_2_ = [early, intermediary, late], we want the total distances between consecutive cell states to be smaller than the distances between non-consecutive cell states. The triangular inequality guarantees that the ratio of these two quantities (the second over the first one) is maximized when the clusters are perfectly aligned in the right order.

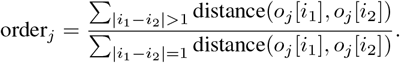
- **Divergence score**: We expect the AML trajectory to diverge from the healthy trajectory. That is, the immature cells of the AML sample are close to the early immature cells of the healthy sample. But then, the blast3 cells of the AML sample are far from the late immature cells of the healthy sample.

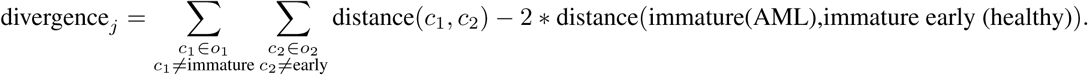

This metric is higher when non-immature AML cells and non-early healthy cells are far from each other and when the immature AML cells and early healthy cells are close to each other.

These metrics attempt to capture our high-level prior knowledge of AML. They summarize the latent space of each method in two numbers: the ordering score and the divergence score. For further details of each method, one can also directly analyze the visualization of the latent space of each method (**Fig. S3**).

Below are the benchmarked methods, the implementation we used, the hyperparameters, and which latent space we used to compute the metrics:

- **PCA**.
  – We run PCA with 50 components (default) using `scanpy`.
  – `sc.tl.pca(adata)`
  – The latent space is the space of 50 PCA components (comparable to our decipher *z* space).
  – `latent = adata.obsm[“X_pca” ]`
- **TSNE**..
  – We run TSNE on the 50-dimensional PCA space using `scanpy` using a knn-graph with *k* = 10.
  – `sc.pp.neighbors(adata, n_neighbors=10); sc.tl.tsne(adata)`
  – The latent space is the 2d TSNE space (comparable to our decipher *v* space).
  – `latent = adata.obsm[“X_tsne” ]`
- **UMAP**
  – We run UMAP on the 50-dimensional PCA space using `scanpy` using a knn-graph with *k* = 10.
  – `sc.pp.neighbors(adata, n_neighbors=10); sc.tl.umap(adata)`
  – The latent space is the 2d UMAP space (comparable to our decipher *v* space).
  – `latent = adata.obsm[“X_umap” ]`
- **Force Atlas**.
  – We run Force Atlas in `scanpy` using a knn-graph with *k* = 10.
  – `sc.pp.neighbors(adata, n_neighbors=10); sc.tl.draw_graph(adata)`
  – The latent space is the 2d force-directed layout space (comparable to our decipher *v* space).
  – `latent = adata.obsm[“X_draw_graph_fa” ]`
- **scVI with batch correction**.
  – We run scVI with two layers, a latent space of dimension 10, and batch correction on the *origin* label (AML vs Healthy), using `scvi-tools`.
  – `scvi.data.setup_anndata(adata, batch_key=“origin”); vae = scvi.model.SCVI(adata, n_layers=2, n_latent=10); vae.train()`
  – The latent space is the 10-dimensional latent space (comparable to our decipher *z* space).
  – `latent = vae.get_latent_representation()`
- **scVI without batch correction**.
  – We run `scVI` with two layers, a latent space of dimension 10, and without batch correction, using `scvi-tools`.
  – `scvi.data.setup_anndata(adata); vae = scvi.model.SCVI(adata, n_layers=2, n_latent=10); vae.train()`
  – The latent space is the 10-dimensional latent space (comparable to our decipher *z* space).
  – `latent = vae.get_latent_representation()`
- **Phate**.
  – We run Phate using the `phate` Python package.
  – `phate_ = phate.PHATE()`
  – The latent space is the 2-dimensional latent space (comparable to our decipher *v* space).
  – `latent = phate_.fit_transform(adata)`
- **Harmony**.
  – We run Harmony using `scanpy` on the PCA with 50 components.
  – `sc.tl.pca(adata); sce.pp.harmony_integrate(adata, ‘origin’)`
  – The latent space is the 50-dimensional PCA-corrected latent space (comparable to our decipher *z* space).
  – `latent = adata.obs[“X_pca_harmony]`
- **Seurat**.
  – We run Seurat using `Seurat` R package.
  – adata <- FindVariableFeatures(data); adata.list <- SplitObject(adata, split.by = “origin”); features <- SelectIntegrationFeatures(object.list = adata.list); adata.anchors <- FindIntegrationAnchors(object.list = adata.list,, anchor.features = features);adata.combin <- IntegrateData(anchorset = adata.anchors); adata.combined <- ScaleData(adata.combined, verbose = FALSE)
  – The latent space is the PCA-corrected latent space (comparable to our decipher *z* space).

### Application of Decipher to PDAC data

We applied Decipher to data collected by Burdziak, et al.[7], consisting of PDAC samples from mouse models with and without KRAS mutation. We subsetted the data to cells undergoing acinar-to-ductal metaplasia (ADM), from three conditions: normal stress, normal, and KRAS-mutated. For our results in **Fig. 3**, we used 10 latent dimensions (*z*), 2 Decipher components (*v*), and *β* = 0.1. All other parameters were default, and the model was run with early stopping. We rotated the resulting Decipher embedding such that the Decipher 1 axis aligned with acinar-to-ductal maturation. Since the desired path of trajectories was previously known, we manually defined trajectories the normal and KRAS-mutated conditions by specifying the order of the clusters.

To interpret the latent dimensions, we selected the latent *z* component that yielded the greatest separation between KRAS-mutated and non-mutated cells. Degree of separation was quantified by T-testing the distribution of a factor over the KRAS-mutated population and the non-mutated population; the absolute value of the T-statistic was used for selecting the best separating component. We also computed the correlation across cells between each latent dimension and each gene **Table 1**. This resulted in a list of genes ranked by correlation values for each latent dimension that may be analyzed either individually, or using Gene Set Enrichment Analysis (GSEA) **Fig. 3e, Table 2**. For individual gene analysis, we looked at genes from the Kras-mutated signature from [7], as well as p53 targets from [11]. To demonstrate that a small set of genes (such as the Kras targets) are ranked significantly higher compared to the ranking distribution of all genes, we applied a Wilcoxon rank-sum test between the set of genes and all genes. Finally, we show the relationship between latent components (z) and Decipher components (v) through visualization the correlation between each *v* and *z* **Fig. 3c**

We repeat the above process using scVI as a comparison. ScVI was run with the same data, using 2 layers and 10 latent layers, with the gene likelihood parameter set to “nb”. The two-dimensional visualization of scVI was obtained using the built-in MDE representation utility **Fig. S1d**. We repeat the same analyses as above on the scVI latent components, identifying the best separating latent component and running GSEA to compare the interpretability of the two methods. We demonstrate that the T-statistics quantifying the degree of separation between KRAS-mutated and non-mutated cells in Decipher’s latent components is significantly higher than in scVI’s latent components (T-test between the Decipher and scVI latent component T-statistics) **Fig. 3g**.

To compare the interpretability of latent factors from the lens of known gene signatures, we utilized the KRAS-mutated signature from [7]. For each Decipher and scVI latent factor, we computed the correlation across cells between each factor and each gene in the signature. We then take the mean of the absolute value of the correlation across genes. We then quantified the separation between KRAS-mutated and non-mutated populations using the T-test, as described above. We plotted the KRAS-signature correlation against the T-statistics to analyze the interpretability of latent factors in each of scVI and Decipher **Fig. 3f, S1f**

### Application of Decipher to AML patient data

Before applying Decipher to the AML patient data, we first performed a gene filtering step to include the most important genes representative of all cell types. For each patient, we performed PhenoGraph [22] clustering using 40 principal components and 30 neighbors. Then, using `scanpy`’s [43] *rank_genes_groups()* function, we performed a T-test to identify the top 400 most differentially expressed genes for each cluster. This list of genes was pooled with a list of known marker genes to produce the final set of genes on which the model was run. We also removed erythrocytes and lymphocytes from the data, as they were not relevant for the analysis of AML derailment.

We obtained:

- 3130 genes for the joint dataset AML1 and normal,
- 3264 genes for the joint dataset AML2 and normal,
- 3258 genes for the joint dataset AML3 and normal,
- 2863 genes for the joint dataset AML13 and normal,
- 2532 genes for the joint dataset AML14 and normal,
- 3291 genes for the joint dataset AML15 and normal,
- 2944 genes for the joint dataset AML16 and normal,
- 2664 genes for the joint dataset AML17 and normal.

Aside from these filtering steps, we emphasize that no other pre-processing or normalization was performed, as the model is always run on raw counts data. For our results **(Fig. 4)**, we use latent factors *z* of dimension 10, Decipher components *v* of dimension 2, a neural network decoder from *v* to *z* with one hidden layer of dimension 64, a linear decoder from *z* to x, a neural network decoder from *x* to *z* with one hidden layer of dimension 128 and another neural network decoder from (*z, x*) to *v* with one hidden layer of dimension 128. BatchNorm was applied after each hidden layer in the neural networks, followed by a ReLU activation. We set *β* = 0.1 and a batch size of 64. The code to reproduce the results in this manuscript is available at https://github.com/azizilab/decipher_reproducibility.

### Interpretation of Decipher components, latent dimensions, and basis

To identify pathways associated with the latent components of Decipher, we computed the covariance of each gene with each of the Decipher components, the latent dimensions, and the results from basis decomposition **(Supplementary Table 2)**. Precisely, we computed for each gene *g* the covariance over cells between *x*_*g*_ and each variable *v*_1_, *v*_2_ and *z*_*j*_ for *j* ∈ [10]. We then ran Gene Set Enrichment Analysis (GSEA) [35], with genes preranked by covariance with each latent component. Next, to interpret the learned basis functions, we ranked genes by their weights in each basis to identify pathways most associated with each basis. For all use cases, GSEA was run on the pre-ranked setting against the Hallmarks Database, with 1000 permutations and no collapse **(Table 4)**. To select genes for visualization in **Fig. S7a**, we selected a top pathway for each component/basis function with known biological importance and found the top disrupted genes belonging to that pathway.

We highlight the usage of Decipher in reconstructing gene patterns over a temporal dimension. These analyses necessitated the translation of cell-level metadata to the temporal dimension. In order to analyze observations such as cell type, mutation proportions, etc., along the temporal dimension, we applied the projection method outlined in the trajectory inference section to obtain a cell-level Decipher time. This transformation allowed for observations to be directly studied along the temporal axis. For discrete observations such as cell type, we first performed nearest-neighbor smoothing using Scikit-learn 0.24.0 [30], with 50 neighbors and a radius of 0.2. A smoothed label was obtained for each cell by taking the mode of the labels among its neighbors. We then visualized the observations along a temporal axis by producing a scatterplot of cell observations, where the x-axis is the computed pseudotime of each cell and the color corresponds to the smoothed label **(Fig. S8)**.

A key feature of Decipher is its ability to produce Decipher components that can be rotated to align with axes of disease maturation and development. We extended our analyses of the NPM1 mutational status of cells to examine its correlation with Decipher component 2 in AML1, 2, and 3. Specifically, we specified a cutoff threshold of 0.4 for the mutation proportion and classified cells with proportions greater than that as belonging to the mutated class and cells with proportions less than as being wild-type. We then binned cells by their pseudotime projections (with a bin size of 0.5 and a sliding window of 0.05). We then counted the number of cells classified into mutated and wild-type by our threshold in each bin and smoothed the resulting counts by time curves using a 1d Gaussian kernel (using `scipy` [40]) with a standard deviation of 2. We visualized the results as distributions along the Decipher component 2 axis to emphasize the shift in NPM1 mutational status **(Fig. 4f, Fig. S3a)**.

### Comparison of Disrupted Genes Across Patients

To determine if disrupted mechanisms between healthy and AML disruption were shared across patients, we first obtained combined disruption scores for each patient as detailed above. We limited our analysis to transcription factors, and identified the top disrupted transcription factors in each patient in order to identify top shared disrupted gene programs. We also visualized disruption scores between patients as a 3D scatterplot of individual patients (Fig. 6d), or by taking the mean of patients with similar mutational statuses (Fig. S11e).

### Distribution of TF peak expression over time

To study the patterns of TF expression over time, we directly utilized the gene patterns produced by Decipher that showed the expression values of each gene over the learned Decipher time axis. For each TF present in the data, expression patterns were first smoothed using a 1d Gaussian filter (using `scipy` [40]) with the standard deviation of the Gaussian kernel set to 3. This smoothing is performed only for detecting peak expression and is not applied to expression plots. Local maxima of expression were then identified by searching for points at which the first derivative of the curve switches from positive to negative. We furthermore filter points by using the midrange of expression (defined as the mean of the minimum and the maximum expression) as a threshold: local maxima whose expression values were less than this value. We additionally included the starting point as a maxima if the expression value was greater than the threshold and the first derivative was negative or if the maximum value of the expression was at the start; we included the ending point as a maxima if the maximum value of expression was at the end. For visualization, we plotted the kernel density estimation (with a bin width of 0.05) of all TF maxima along the Decipher time axis to show the points in time where overall TF activity is concentrated **(Fig. S8)**. Kernel density estimation was performed using `seaborn` [42], and plotting was done using `matplotlib` [16].

### Analysis of temporal TF co-regulation

In conjunction with the visualization of the timing of TF activity, we also sought to determine if families of similar TFs demonstrated coordinated activation times and if these temporal dynamics could be utilized to derive insight into regulatory wiring. We focused our analyses on the top 20 most disrupted TFs by the combined disruption metric, as well as the known disrupted TFs from the literature. The Decipher gene pattern for each TF was visualized as rows in a heatmap, with the horizontal axis representing the pseudotime axis and the color representing the z-scored expression value. The rows were sorted based on the time at which the maximum peak occurred, and the TFs were labeled with colors based on their biological function (**Fig. 6b,c; Fig. S10a,b**).

### Estimation of uncertainty in expression patterns

Because Decipher is a probabilistic model, it learns the uncertainty about the gene expression induced by a cell representation *v*. Given a location *v* in the Decipher space, the distribution of the expected gene expression *µ*_*g*_(*v*) of gene *g* in a cell with representation *v* is given by,

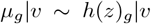

where *z*|*v ∼ 𝒩* (*f*_mean_(*v*), *f*_var_(*v*)).

To compute this uncertainty for each *v*, we sample 100 values for *z* from *z* | *v* ∼ 𝒩(*f*_mean_(*v*), *f*_var_(*v*)) and compute *µ*_*g*_ = *h*(*z*)_*g*_ for each of them. In **Fig. 5b**, the shaded bands represent the interquartile range (25-75) of the 100 samples.

### Analysis of bulk AML data

We applied DeSeq2 [26] to obtain metrics characterizing the AML data at the cohort level. Using the resulting L2FC and p-values, we were able to confirm whether or not expected genes, as well as our newly identified disrupted genes, were also differentially detected in the bulk data. For both sets of genes, we selected genes whose absolute L2FC was greater than 1 and reported the maximum p-value.

### Application to gastric cancer evolution

We applied Decipher on the gastric cancer data from Kim *et al*. [19]. *We pooled the data from the 24 patients in the study, each with pre-malignant cells and cancerous cells. 9 of these patients have intestinal* cancer and the other *15* patients have *diffuse-like* cancer. The resulting data has 12,612 cells and 8,705 genes. We ran Decipher with its default hyperparameters: *z* of dimension 10, *v* of dimension 2, and for 30 epochs **(Fig. 7b-d)**.

In sum, Decipher on the gastric data was applied on

- 12614 cells,
- 8705 genes,
- from 24 patients,
- with 2 major types of cells (pre-malignant, cancerous)
- with 2 types of cancer (diffuse-like or intestinal),
- with different stages of cancer.

## SUPPLEMENTARY INFORMATION

### Semi-synthetic simulation

To further complement our validation of Decipher’s performances, we created semi-synthetic data that is more realistic than purely simulated data, by leveraging the AML data. We used the Decipher model trained on real data (normal HSPCs and sorted cells from AML1) to obtain realistic values for the probabilistic model. Then we simulated new data by (i) forming two divergent ground-truth trajectories in the Decipher component space (Methods), (ii) generating synthetic cells by sampling their Decipher components uniformly along these trajectories, and (iii) generating synthetic gene expression values for these simulated cells using our probabilistic model (**Fig. 1d; Fig. SI1a,b**). Then, we fitted a new Decipher instance to this simulated dataset that is realistic and for which we have ground truth. Decipher successfully learned a latent space in which cell-state transitions from immature to blast0–3, and both normal and AML trajectories are correctly aligned along the Decipher 1 component (**Fig. SI1c**). The derailment in the AML condition is captured by the Decipher 2 component, which precisely locates the bifurcation point. We assessed the recovery of cell order by evaluating the correlation between the inferred Decipher time and the ground-truth trajectory. To confirm the robustness of these results, we repeated the entire Decipher pipeline across five simulated datasets and consistently observed a correlation above 0.94. Finally, we examined robustness in the context of rare or poorly represented cell states, similar to the fully synthetic experiments. We removed 90% of cells from two intermediary cell states (**Fig. SI1c**) and fitted Decipher on this new simulated data. Even with a non-uniform sampling of cell states, Decipher successfully preserved the order of cell states (**Fig. SI1d**), as found in the fully synthetic experiments.

**Figure S1.**
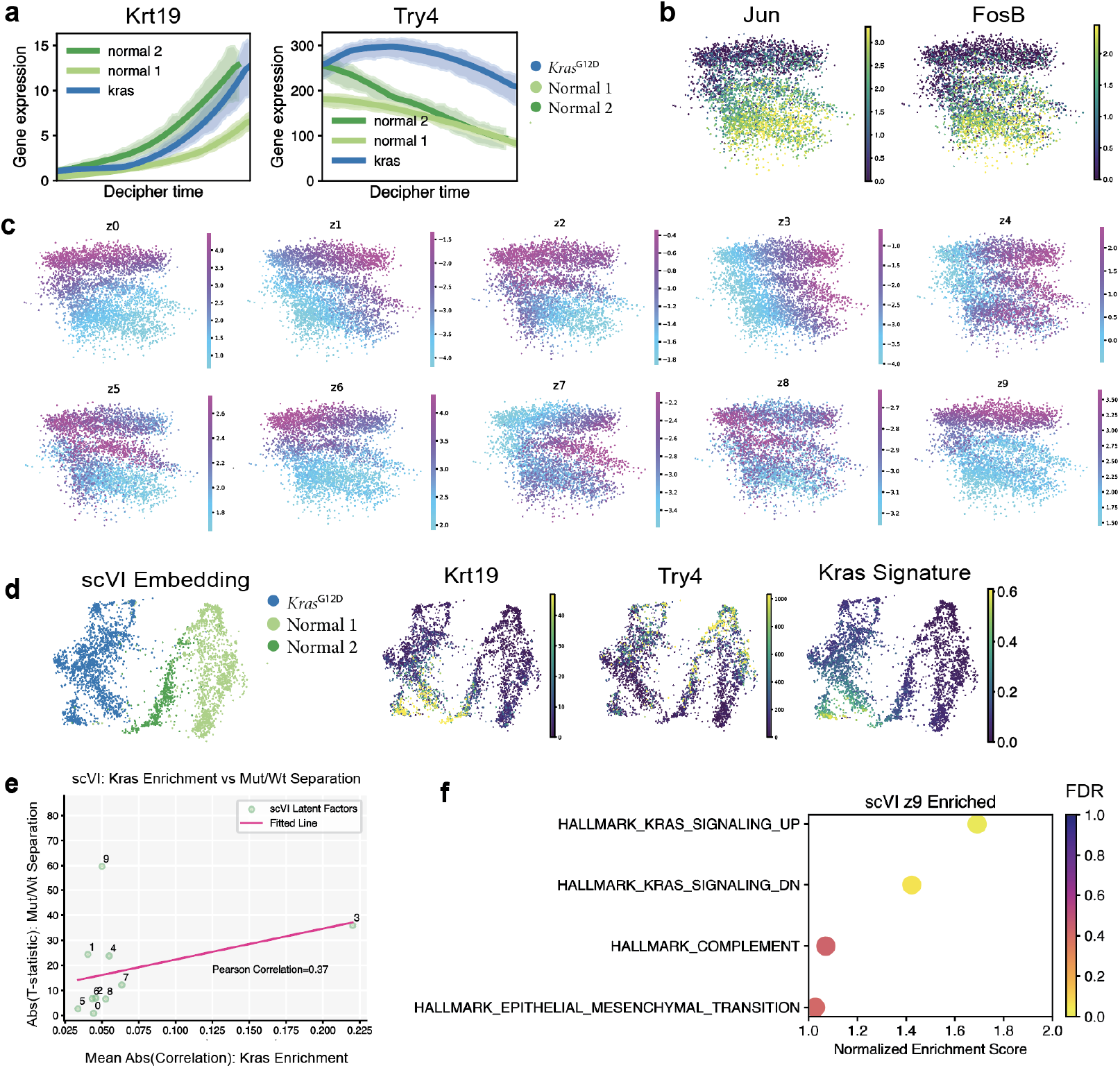
Comparison of latent spaces of scVI and decipher. **(a)** Expression of ductal marker *Krt19* and acinar marker *Try4* over learned Decipher time. **(b)** Decipher embedding colored by AP-1 factor genes *JUN* or *FOSB*. **(c)** Decipher embedding colored by latent factors. **(d)** 2D minimum distortion embedding (MDE) computed from scVI latent factors, colored by mutational status, gene markers, and the *Kras*-mutated signature. **(e)** Correlation between the absolute value of the t-statistic quantifying the distance between *Kras*-mutant and normal cells in each scVI latent factor, and the mean absolute value of each factor’s enrichment in the bulk *Kras* mutational signature. **(f)** Pathways enriched by GSEA for z9 in scVI, which is the factor with the biggest separation of *Kras*-mutant and normal cells, show *Kras* signaling both up- and downregulated.

**Figure S2.**
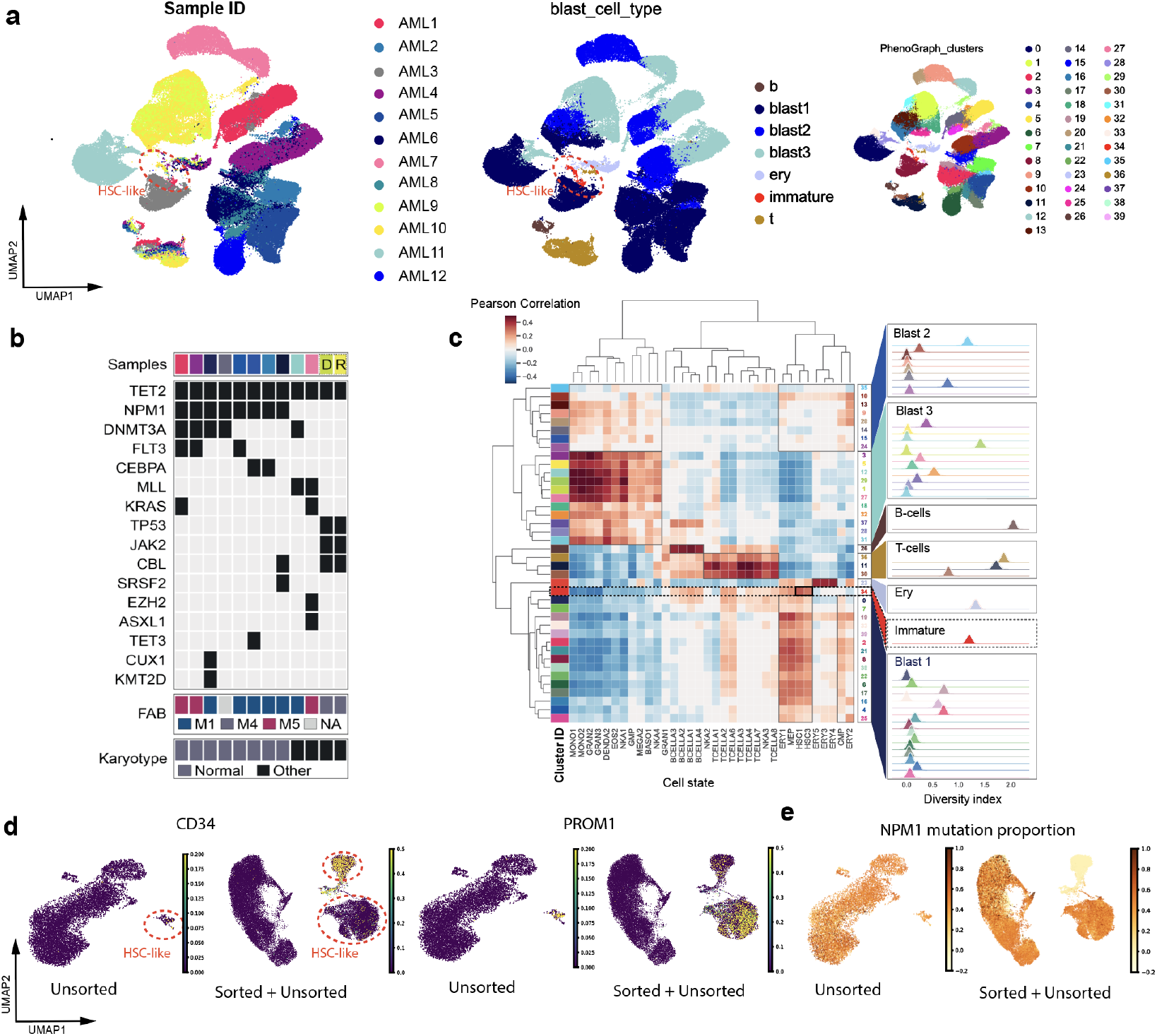
Characterization of AML samples and immature cell enrichment. **(a)** UMAP projection of unsorted bone marrow transcriptomes from 11 AML patients (AML9 and AML10 are from the same patient). Cell colored by patient ID, cluster ID, or annotated cell type. **(b)** Targeted mutation, karyotype, and morphological (FAB) assessments of patient bone marrow samples (**Table 3**). D, R denote diagnosis and relapse paired samples, respectively, from the same patient. **(c)** Left, Pearson correlations between cluster centroid expression and bulk gene expression data from sorted subsets of healthy HSPCs[64] (Methods). Right, Shannon Diversity index computed for the distribution of patients in each cluster, controlling for cluster size (Methods). Higher diversity indicates greater mixing of patients. **(d)** UMAP projection of AML1 single-cell transcriptomes including or not including sorted CD34^+^/PROM1^+^ cells. Cells are colored by the expression of CD34 and PROM1 and **(e)** proportion of *NPM1* mutation in their neighborhood (Methods).

**Figure S3.**
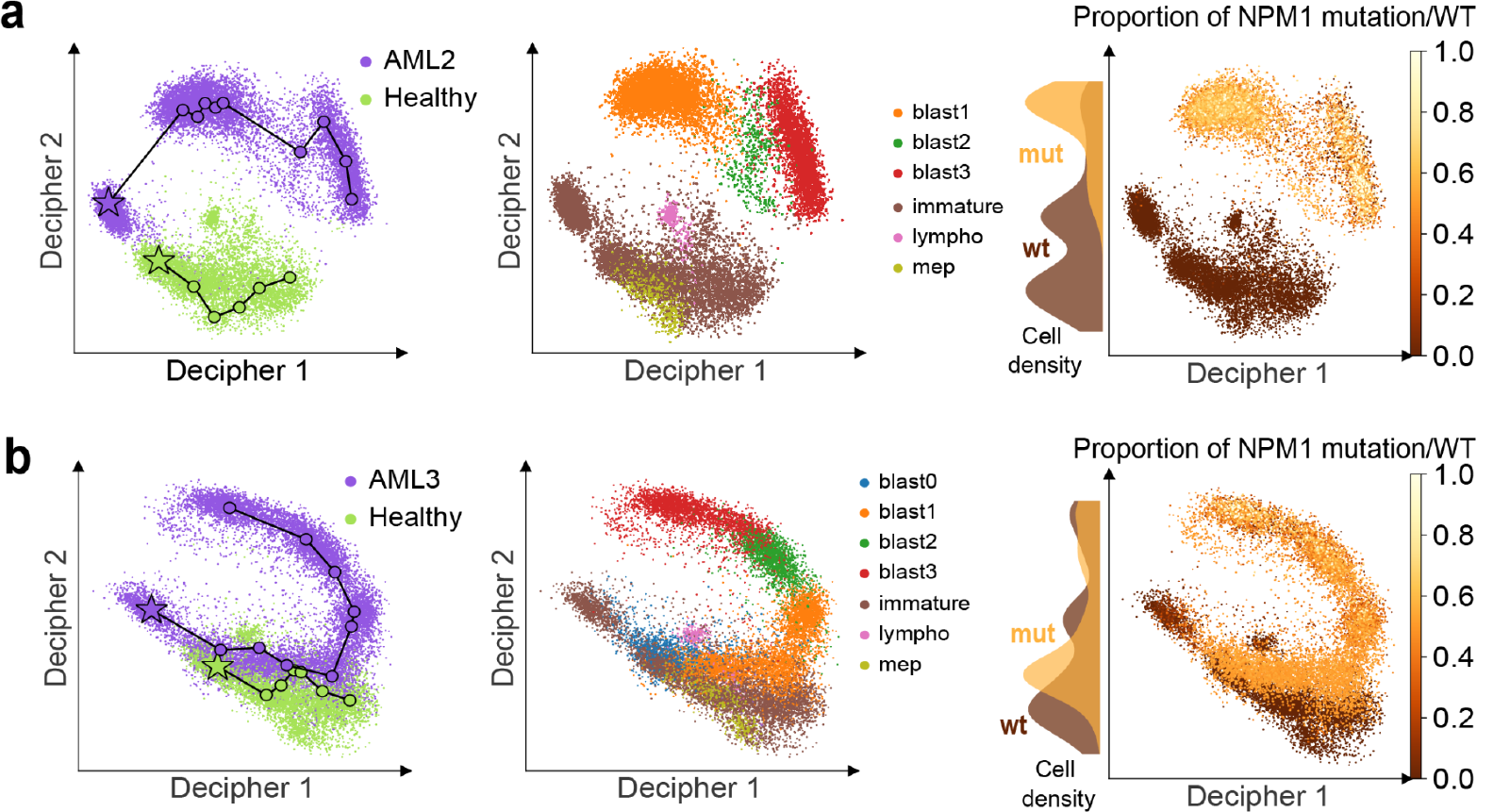
Application of Decipher to other AML patients. **(a,b)** Decipher space visualization of cells from the healthy donor and AML2 **(a)** or AML3 **(b)**. Cells colored by condition (left), cell type (middle), and proportion of *NPM1* mutation (defined as the number of neighbors with mutation/all neighbors for each cell, using 30 neighbors) (right). Lines define trajectories for normal and healthy samples (left).

**Figure S4.**
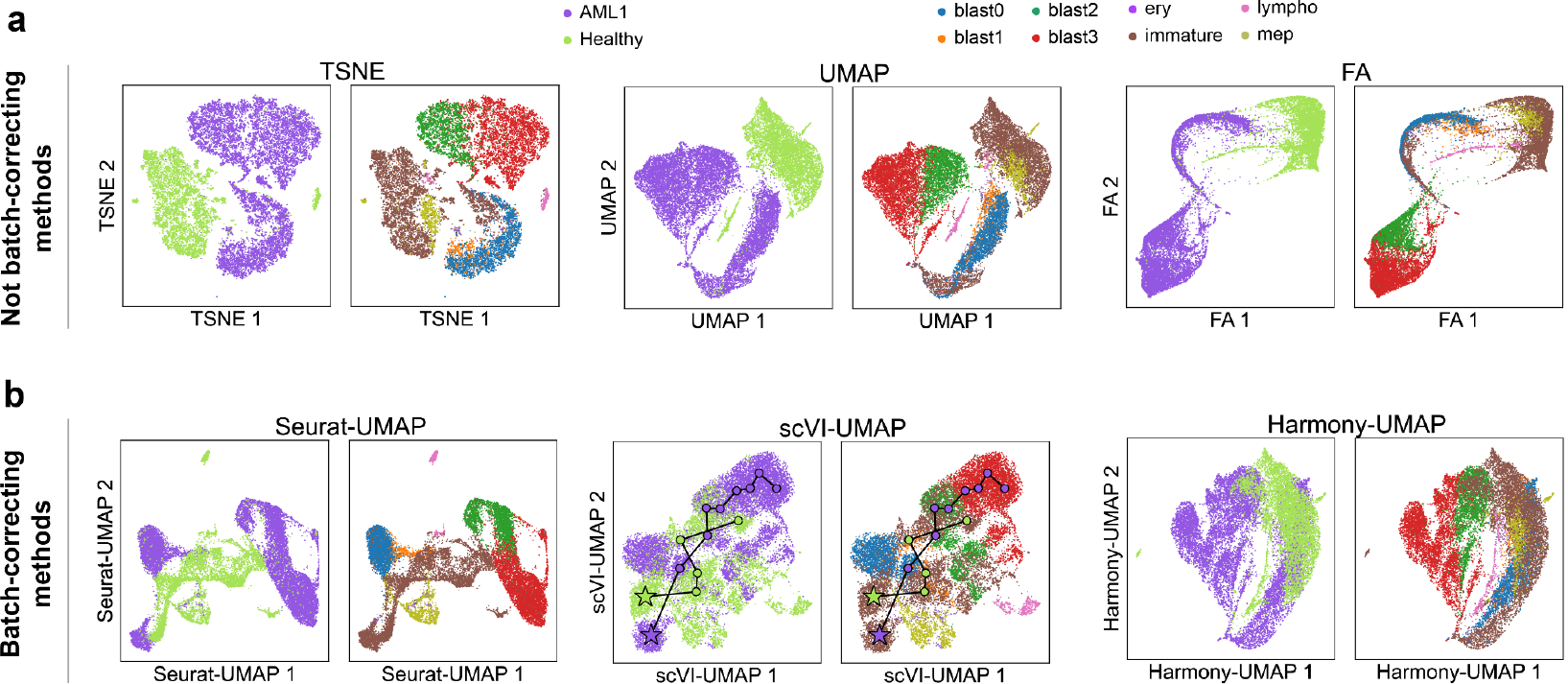
Integration of AML and normal samples with existing methods. **(a,b)** Projection of 37,395 cells from patient AML1 and from a healthy donor using different tools (Methods). Each dot represents a cell colored by origin (left) and type (right). Non-batch-correcting visualization methods tSNE, UMAP, and FDL (a), and batch-correcting methods – UMAP of the Seurat space, scVI space, and Harmony space (b) – all fail to integrate AML and healthy cells. Lines indicate trajectories paths inferred on the scVI space (Methods).

**Figure S5.**
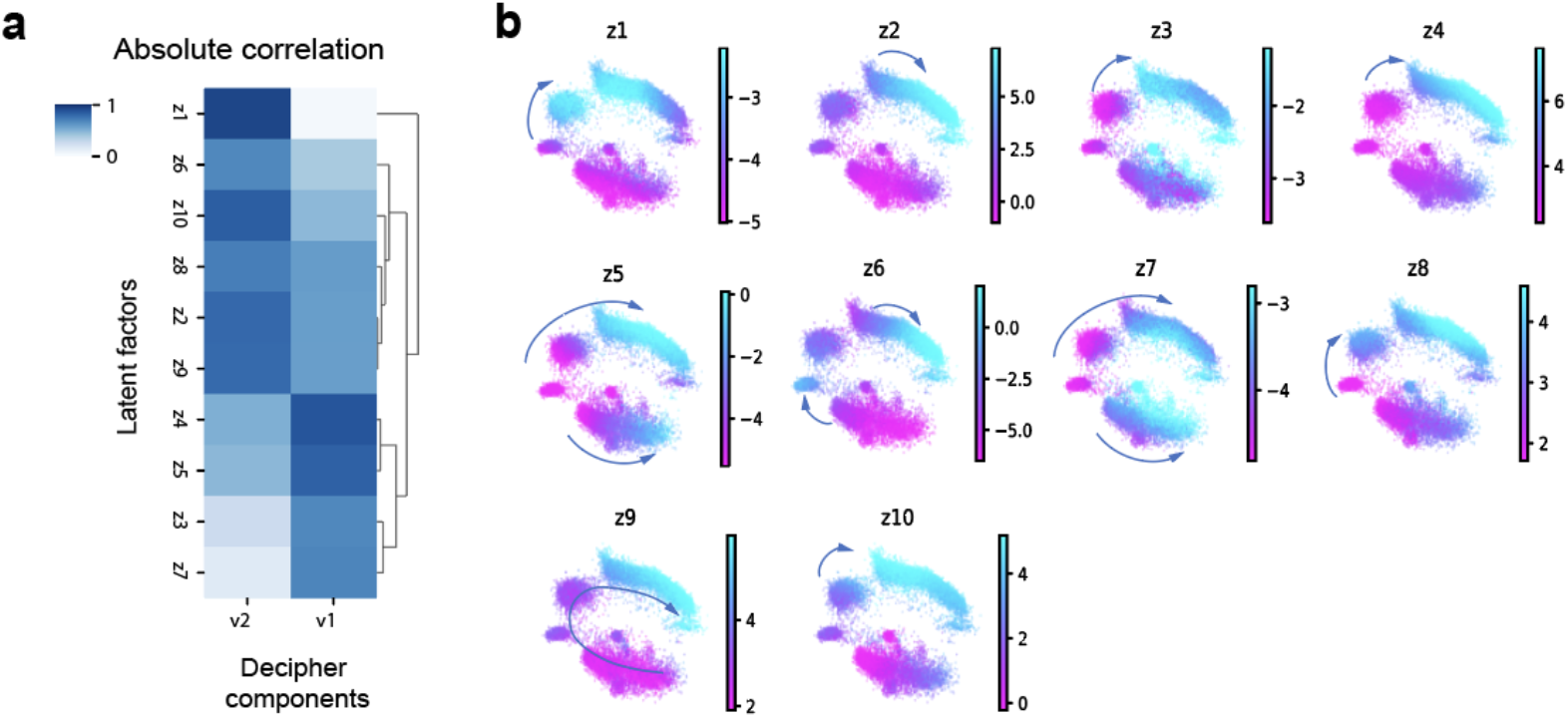
Latent factors capture AML cell-state transitions. **(a)** Absolute value of correlation between Decipher components and latent factors. **(b)** Decipher space colored by all latent factors for AML1 (a subset of these are shown in **Fig. 4h**). Arrows indicate cell-state transitions corresponding to an increase in the latent factor.

**Figure S6.**
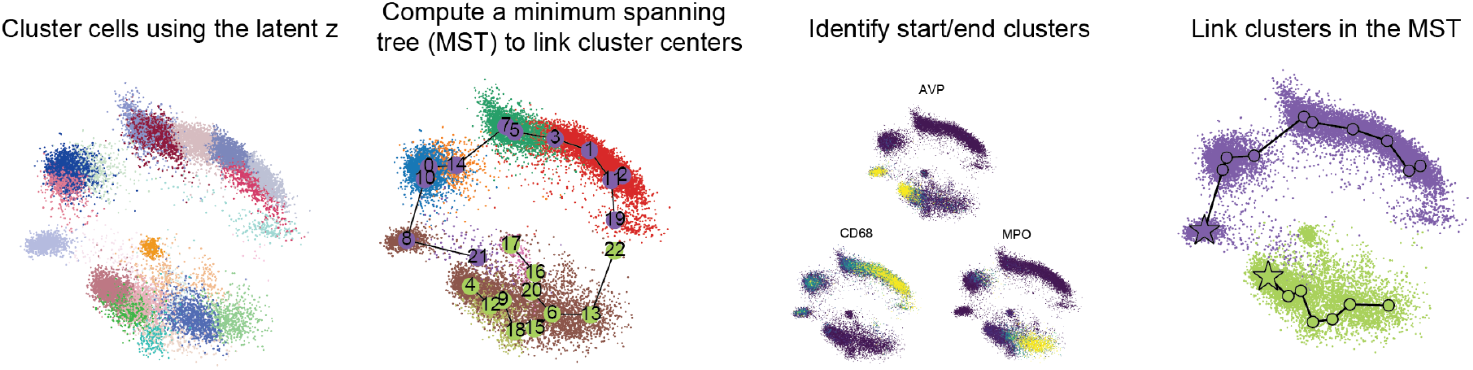
Decipher trajectory inference. Steps involved in trajectory inference, from left to right, based on the inferred representation from Decipher (Methods).

**Figure S7.**
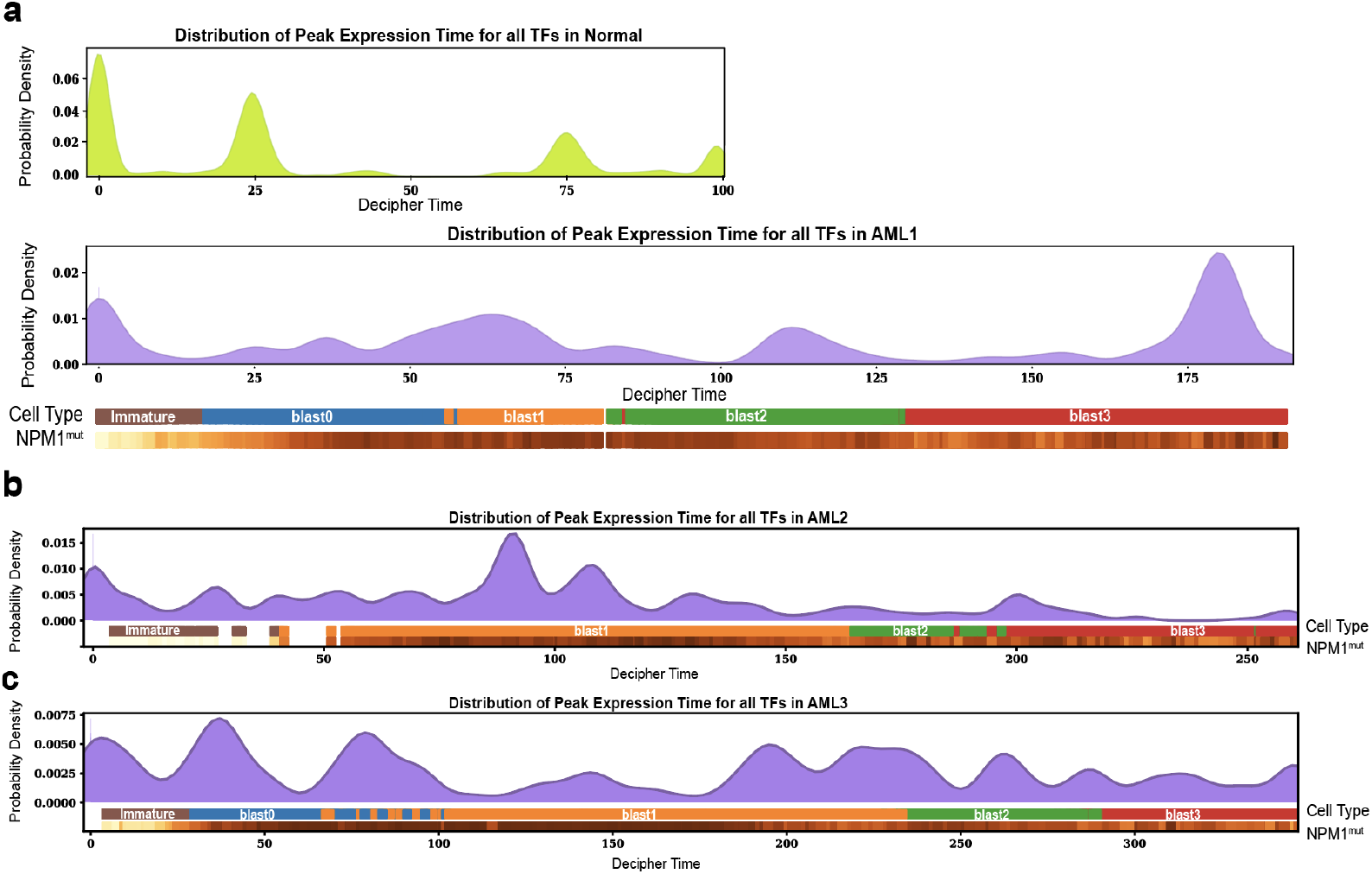
Temporal distribution of genes and TFs. **(a)** Timing of peak expression of TFs on the Decipher time axis in normal (top) and AML1 (bottom) samples. Density plots display the local maxima across all TFs in each sample. Distribution of local density maxima across all TFs along Decipher time in AML2 **(b)** and AML3 **(c)**.

**Figure S8.**
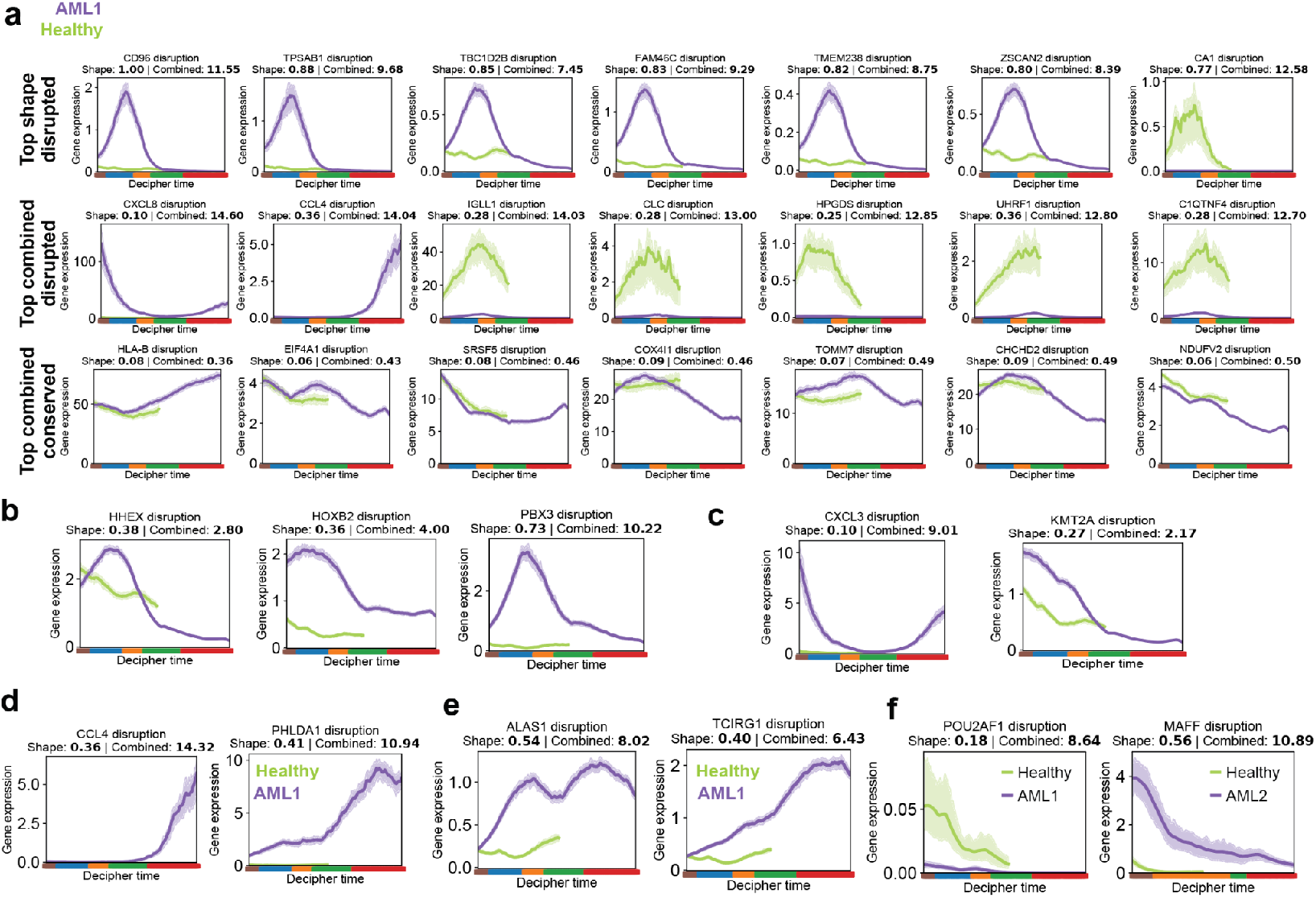
Gene trend analysis using Decipher. **(a)** Reconstructed expression patterns for top disrupted/conserved genes that were identified using the shape and combined disruption metrics. **(b–e)** Reconstructed expression trends and disruption scores for homeobox genes and cofactors (b), genes upregulated in immature leukemic cells (c), TNFɑ pathway (d), oxidative phosphorylation (e) in AML1. **(f)** Reconstructed trends and disruption scores for genes disrupted mainly in early immature state and thus unidentified in bulk analysis.

**Figure S9.**
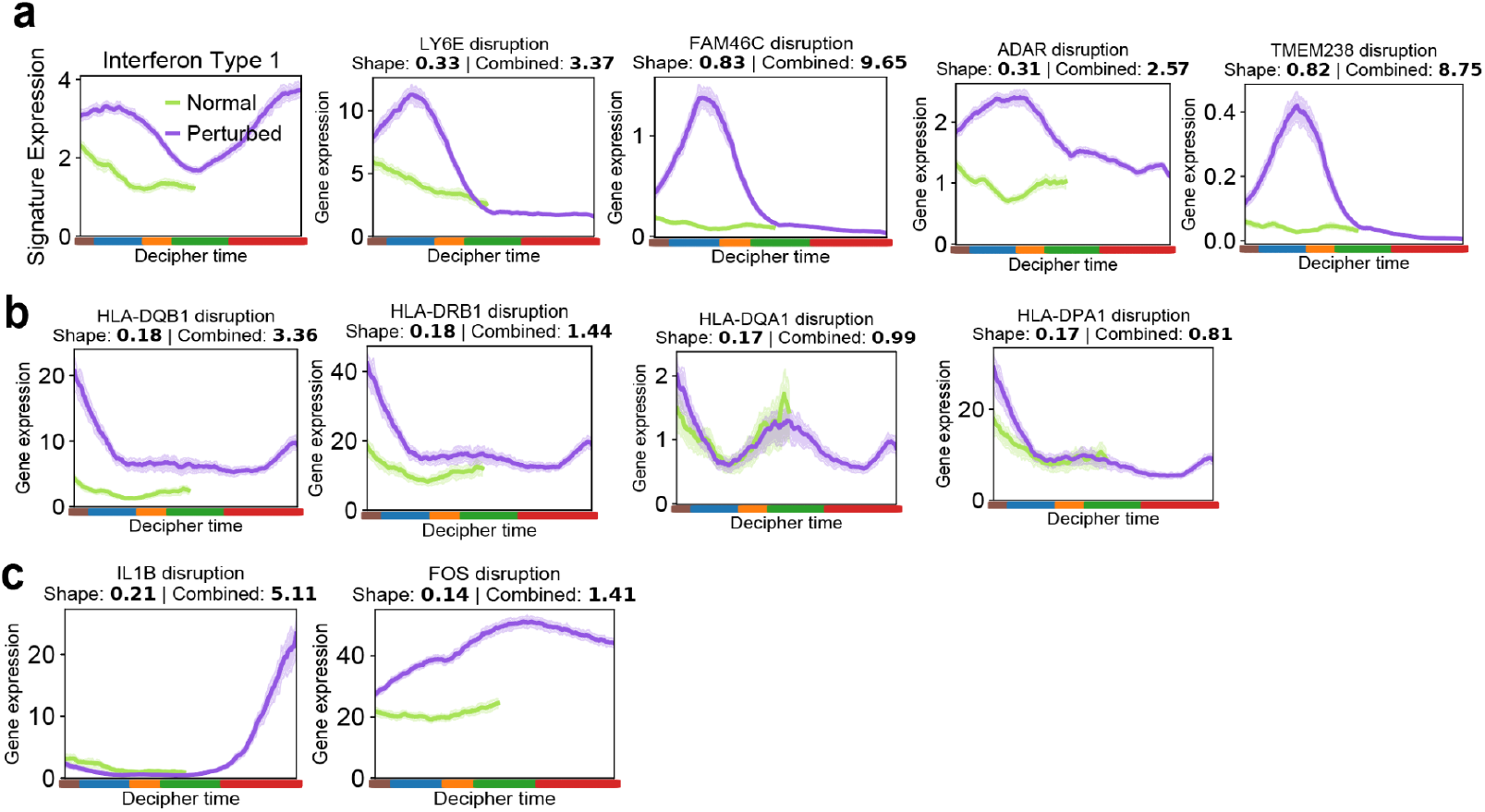
Notable genes co-expressed with aberrant TFs and comparison of TFs across patients. **(a)** Reconstructed expression trends for IFN type 1 signature (left) and individual genes (right) found to be co-expressed with *HOX* TFs under both normal (green) and AML (purple) conditions. **(b,c)** Reconstructed expression of genes involved in IFN type 2 (b) and TNFɑ pathway (c).

**Figure S10.**
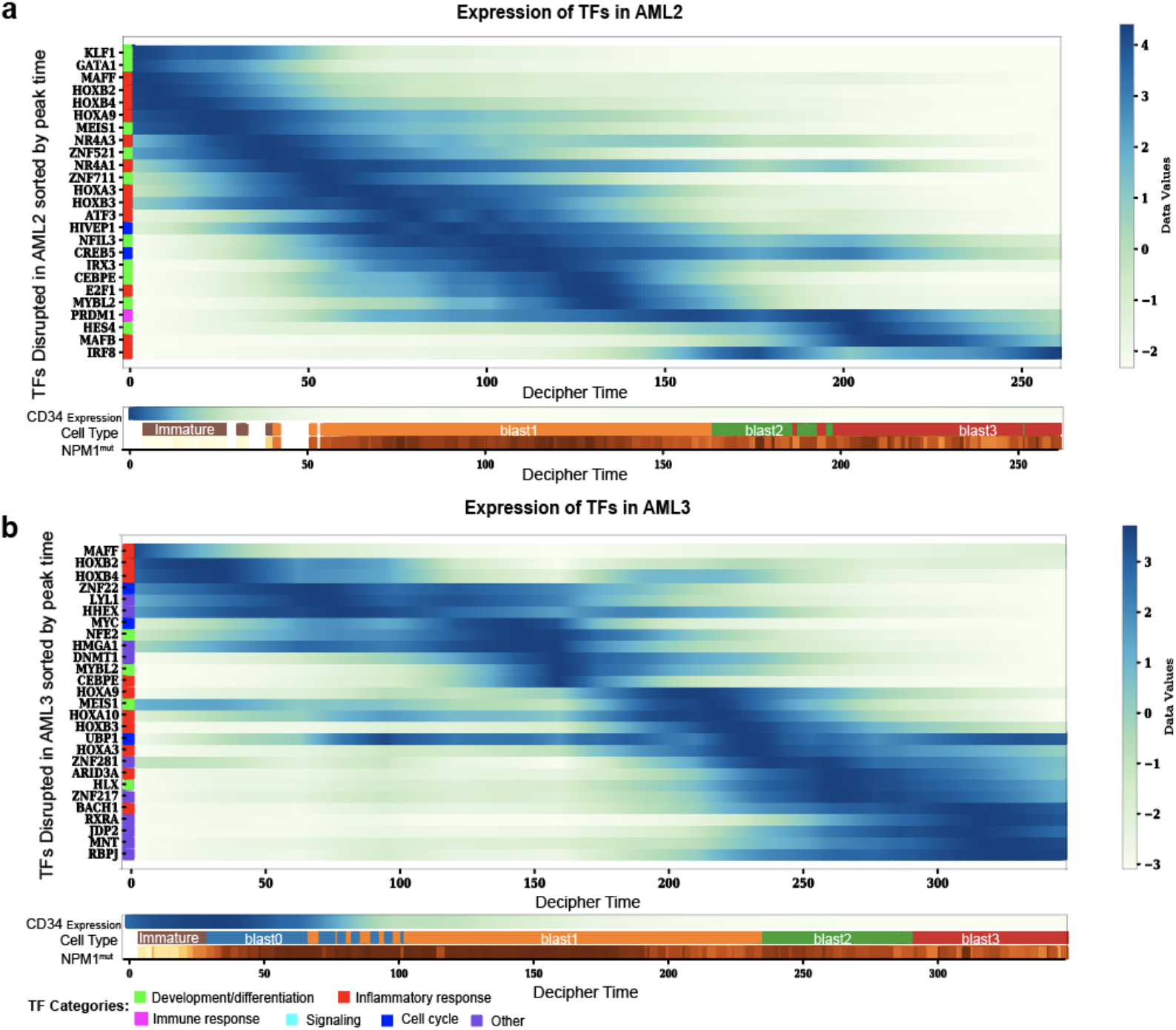
Altered TF dynamics in patients AML2 and AML3. **(a,b)** Timing of TF expression along Decipher time in AML2 (a) and AML3 (b) samples. Heatmaps show log-transformed and z-scored expressions for the top 20 TFs with the highest combined disruption scores in each sample, as well as known TFs from literature, sorted by timing of peak expression. Colorbars correspond to cell type and proportion of *NPM1*-mutated cells among the 30 nearest neighbors of each cell; both are smoothed over the 50 nearest neighbors in the Decipher space.

**Figure S11.**
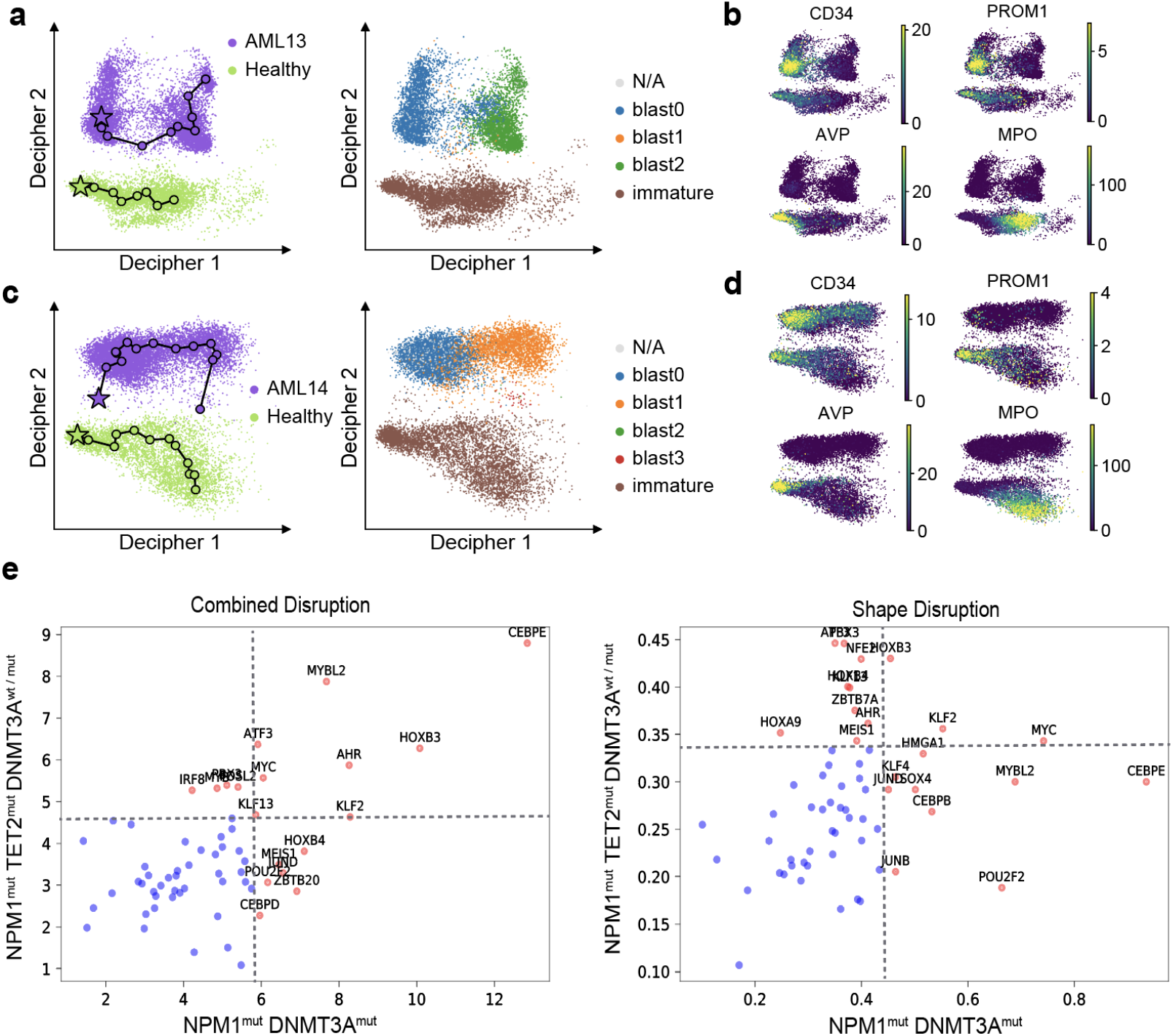
Decipher characterizes a *DNMT3A*-mutated AML cohort. **(a–d)** Decipher space embedding of sorted cells from two *DNMT3A*^mut^ AML patients (Table 3), colored by sample of origin and cell type (a,c) and by marker gene expression (b,d). AML maturation is observed as a decrease in *CD34* expression, and normal HSPC differentiation as a decrease in *AVP* and increase in *MPO*. Decipher identifies and aligns myeloid trajectories in AML even when cell states are missing (blast1,3 in AML13; blast2,3 in AML14). **(e)** Comparison of disrupted genes in patients bearing *NPM1* and *DNMT1* mutations, with and without *TET2* mutation, based on combined disruption and shape disruption scores. Red points represent TFs in the 80th percentile of disruption in at least one category. The dotted lines show the 80th percentile threshold.

**Figure SI1.**
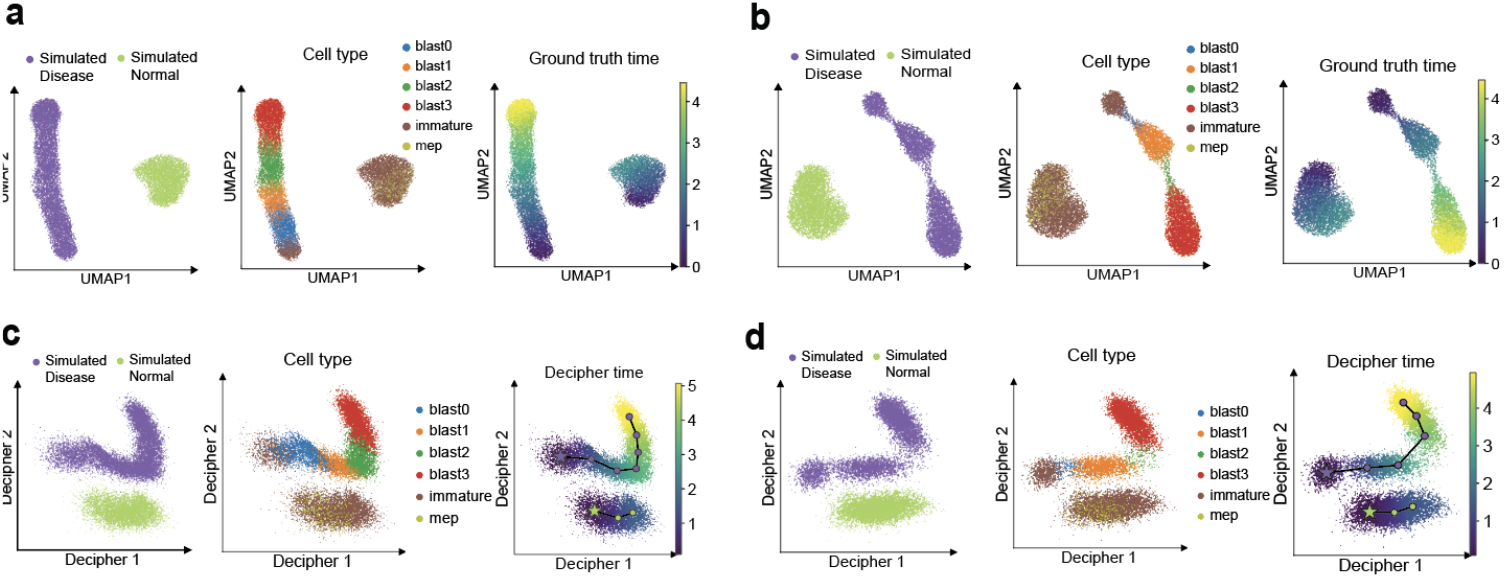
**(a)** UMAP visualization of semi-synthetic data (18,000 cells) simulating an AML patient and a healthy reference (Methods) along simulated trajectories. Each dot represents a cell colored by sample (left), cell type (middle), and ground truth time (right). **(b)** UMAP visualization of the data simulating rare cell types. Similar to (a) but removing 90% of blast0 (blue) and blast2 (green). Each dot represents a cell colored by sample (left), cell type (middle), and ground truth time (right). **(c)** Example of Decipher applied to the data in (a). It preserves the order of cell states and shows the shared maturation (Decipher1) as well as the divergence of mature leukemic blast cells (blast2,3) in AML (Decipher2). Each dot represents a cell colored by sample (left), cell type (middle), and inferred Decipher time (right). **(d)** Decipher applied to data shown in (b) can still learn meaningful representation and trajectories.

### Design of AML Experiment

To characterize the impact of NPM1mut while minimizing confounding effects, we focused on patients bearing TET2 epigenetic mutations (n = 12), with and without NPM1 mutations. We used scRNA-seq to profile 104,116 AML bone marrow cells (**Fig. S2a** and **Table 3**). Assuming that leukemic cells preserve a subset of normal hematopoietic differentiation programs, we used well-characterized normal references [64] to achieve cluster-level annotation of the maturation stage (**Fig. S2b**; Methods). This analysis revealed a broad spectrum of leukemic cell states that partially recapitulates the diverse morphological subtypes in our cohort [99,100] (**Fig. S2c**); differentiated (M1) samples were mostly populated by immature blasts, myelomonocytic (M4) and monoblastic (M5) samples by cells with greater maturation. However, most samples contained many cell types.

Comparison of pre-leukemic cell and normal HSC phenotypes consistently returned PROM1 as a marker for the immature population in human NPM1mut AML samples. This analysis also identified consistent underexpression of HSC genes such as CD34 and AVP in NPM1mut cells (**Fig. S2d,e**), concordant with the demonstrated ability for NPM1 mutations to induce HOX genes in vitro [101] and confer self-renewal properties to myeloid progenitors in mice [59]. Overall, this approach provides an expanded map of both frequent and rare cellular populations that exist in each patient and unambiguously delineates NPM1 mutation status.

### Analysis of single-cell ATAC-seq

To further study altered TF activity in AML, we performed an integrated analysis of gene expression and chromatin accessibility patterns via single-cell Assay for Transposase-Accessible Chromatin (scATAC-seq; Methods). Using scATAC data, we explored possible regulatory relationships between disrupted TFs expressed in early immature cells and those expressed later in AML progression that could explain the observed cascades of TF activity (Fig. 6d; Fig. SI2). Specifically, we identified the top 50 disrupted TFs for each patient, and split them into those expressed before and after NPM1 mutation, labeled as ‘early TFs’ and ‘late TFs’ respectively. We then investigated putative regulatory relationships between the two groups of TFs by computing a motif enrichment score for each early TF detected within accessible regions in the vicinity of late TFs as targets. Then, using bootstrapping analysis, we compared the motif scores to a null distribution generated from same-size random sets of target TFs that are expressed in at least 1% of cells (Methods). In all three patients, we found significant motif enrichment in HOX TFs, including HOXB4 (p < 0.0458) and HOXB6 (p < 0.032), and FOSL1 (p < 0.0057), suggesting their putative role in regulating TFs driving AML transformation. In two out of three patients, we found significant motif enrichments in GATA2, MYC, NR4A2, MAFF, ATF3, NFKB2, RELB, KLF2, ZNF274 (p < 0.05; Methods).

We then examined the dynamic accessibility of the TFs themselves during AML progression through the integration of scRNA- and scATAC-seq data using LIGER[36] (Fig. SI3a,b, Methods). Through this process, we are able to project scATAC data into the 2D Decipher space inferred from scRNA (**Fig. SI2c**), finding that the patterns of chromatin accessibility mirror gene expression dynamics for regulators with enriched motifs (**Fig. SI2d**). We find that the significantly enriched motifs (GATA2, MYC, NR4A2, MAFF, ATF3, NFKB2, RELB, KLF2, ZNF274) show altered expression in immature cells in two out of three patients (**Fig. SI3**).

Interestingly, motif enrichment analysis using scATAC data further showed enrichment of motifs for known interferon-induced TFs such as STAT1 (MA0137.1,p<0.0357) and IRF1 (MA0050.1,p<0.0057; MA0050.2,p<0.0216) among the top disrupted TFs in AML1 and AML3. Our approach thus resolves the timing of TF activity with respect to significant events such as genetic mutations and activated signaling pathways, guiding further studies of regulatory relationships.

**Figure SI2.**
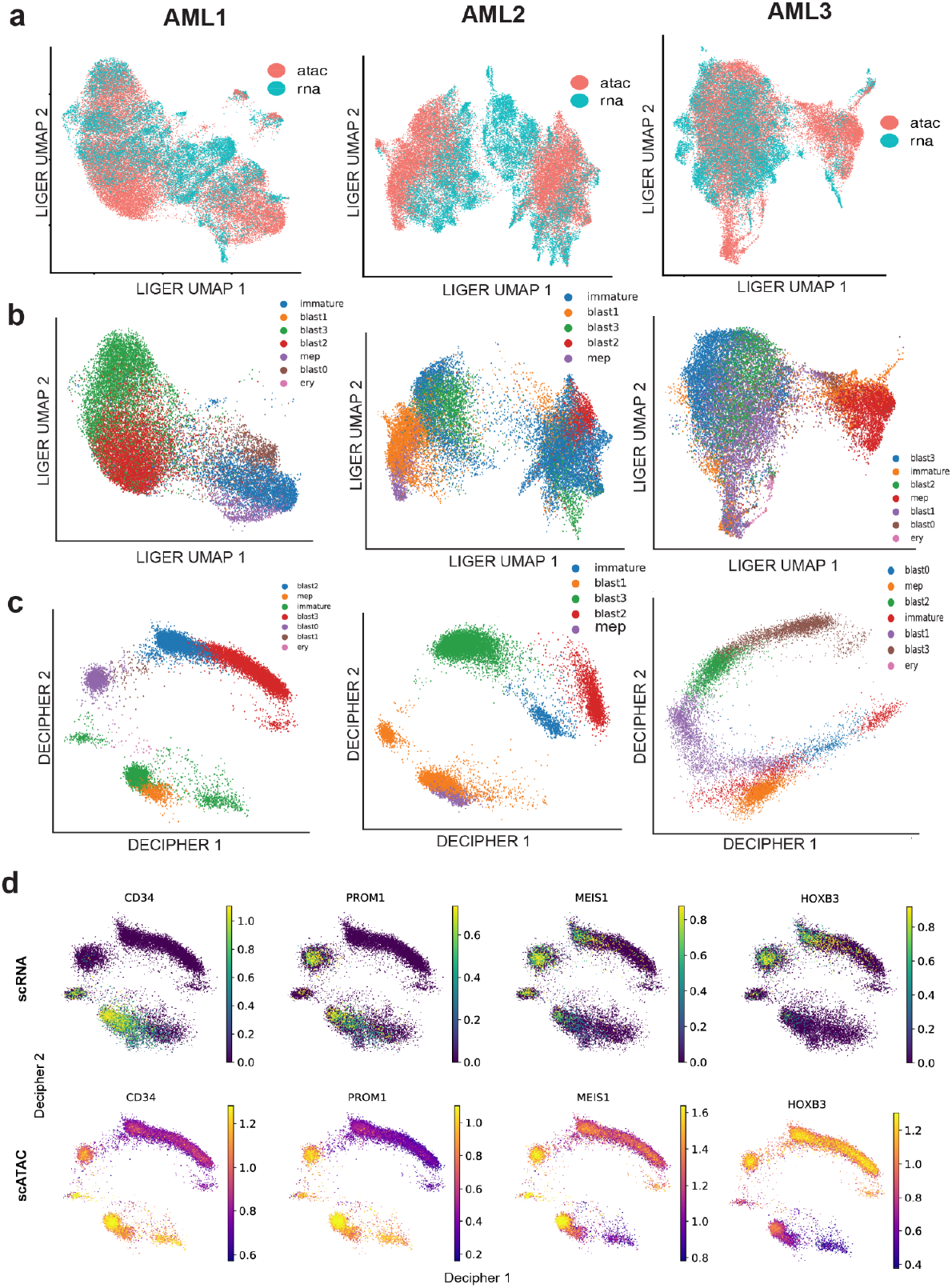
Integration of scRNA and scATAC data. **(a)** UMAP embedding of cells from scATAC and scRNA-seq in the LIGER integrated space in three AML patients. **(b)** UMAP projection of cells colored by cell types in ATAC-seq in the LIGER Integrated space in three AML patients. **(c)** Projection of cells in scATAC data in 2D decipher space colored by cell type. **(d)** scRNA and scATAC-seq of key marker genes in AML1. Scatterplots show expression (top) and accessibility (bottom) in the Decipher space (Methods).

**Figure SI3.**
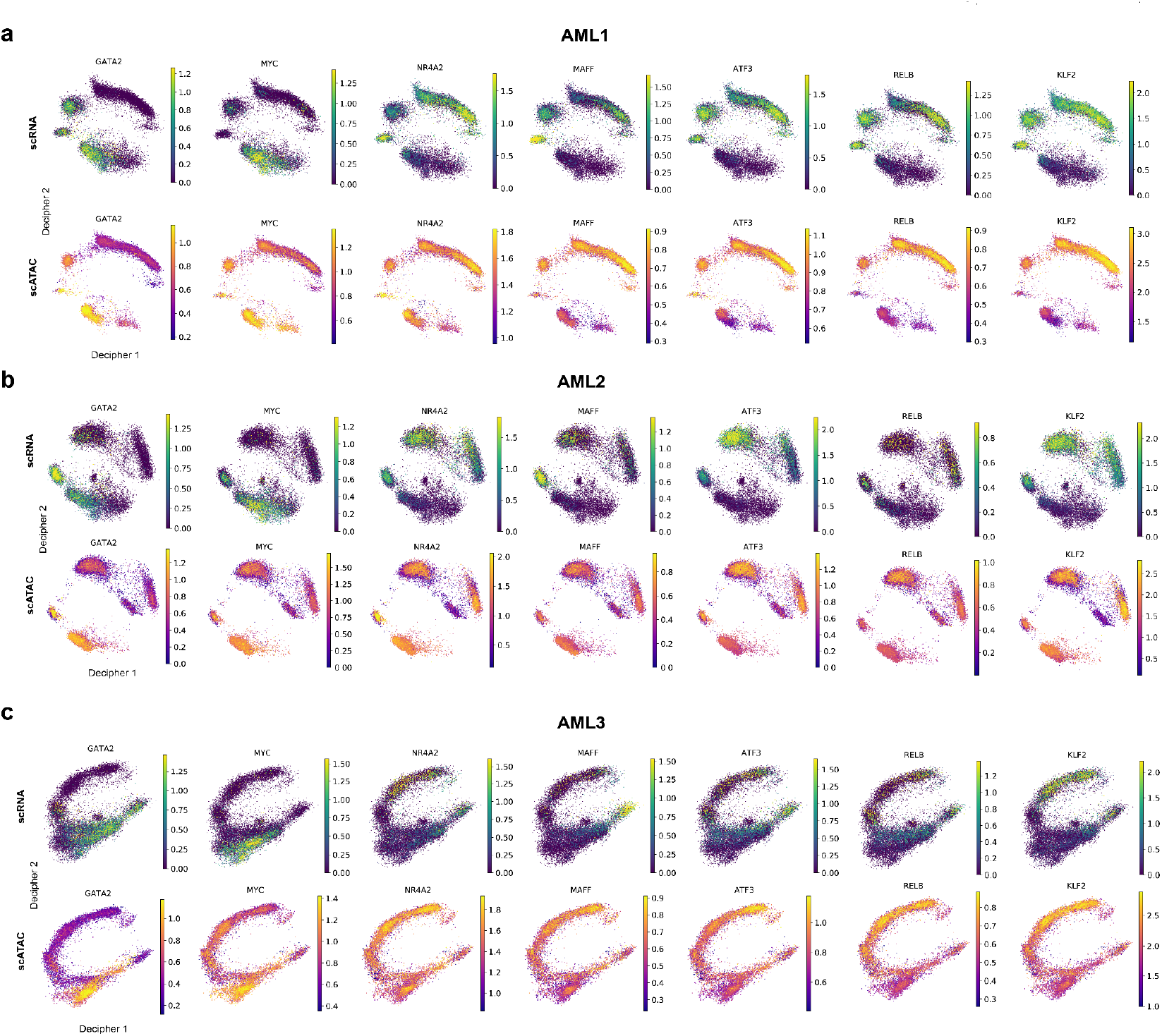
TFs with enriched motifs in all patients. scRNA and scATAC-seq of TFs found to have enriched motifs in the vicinity of late TFs (expressed after NPM1 mutation) in two out of three AML patients (Methods). Scatterplots show expression (top) and accessibility (bottom) in the Decipher space for AML1 **(a)**, AML2 **(b)**, and AML3 **(c)**.

### Analysis of single-cell ATAC-seq

We utilized ArchR[102] for pre-processing of scATAC-seq data. In order to analyze accessibility trajectories from immature cell states to mature blast cells, we combined each patient AML sample with scATAC-seq data collected from a healthy bone marrow sample. ArchR was run using default parameters and the HG38 genome for peak alignment. UMAP visualization was run using ArchR’s implementation with iterative LSI in order to provide a preliminary visualization of the distribution of cells in the ATAC space. The peak counts and gene scores were then exported and stored in an AnnData object for downstream use in *Scanpy*.

### Integration of scATAC and scRNA

To integrate the scATAC data with the scRNA data, we applied LIGER[36], using the gene scores from ArchR and the counts from the scRNA data as the two inputs. We applied a log2(*x*+1) transformation to the gene scores (where x=gene scores) to improve the signal. Running with standard parameters yielded an integrated space that could be visualized using UMAP (**Fig. SI2a,b**) to confirm the overlapping of modalities and improved separation of cell types. From this step, we exported the matrix of normalized cell factors, *H*, that represents the LIGER shared factors across modalities that can be subsequently analyzed to compare cells from the scATAC data to cells in the scRNA data.

### Projection of Accessibility scores in Decipher space

The first step of this analysis was to project cells in scATAC-seq to the Decipher space by finding the most similar cells in the RNA-seq to each cell in the ATAC-seq, with the goal of obtaining sharing metadata and Decipher values from the RNA-seq to the ATAC-seq. To achieve this, we applied a nearest neighbors algorithm using Scikit-learn [103] to accomplish the projection as follows: we first trained a neighbors classifier on the *H* matrix obtained from LIGER for RNA alone, *H*_*RNA*_, (with the number of neighbors set to 3). Then, for each cell in scATAC-seq, we applied the classifier to the corresponding row *H*_*ATAC*_. The classifier subsequently returned the 3 nearest cells in RNA for each cell in ATAC-seq. We then compute the projections by taking the mean Decipher *v* component value for each cell, as well as the mean projected value onto the trajectory (Decipher time). The annotation for a given cell was simply taken to be the mode cell type of the neighbors (**Fig. SI2c**).

Because this projection step may be a noisy process, we also implemented a filtering step wherein we computed the maximum pairwise Euclidean distance using *Scipy [104]* of the Decipher space coordinates in each set of 3 nearest neighbors. The cells with distances greater than a selected threshold were removed. This step was particularly important for those Decipher embeddings that take on a curved or horseshoe shape, where a cell from ATAC-seq may be assigned to cells originating from two opposing regions.

Finally, to obtain a better signal for visualizations, we again implemented a nearest-neighbors smoothing. We first built a nearest neighbors graph (k=50) on the ATAC peaks matrix. Then, the accessibility values for a given cell are obtained by taking the mean gene score for each gene across all neighbors (**Fig. SI2d**)

### Comparison of scRNA-seq and scATAC-seq in the Decipher space

For key transcription factors and marker genes, we sought to compare the patterns in their expression and their accessibility. For the expression plots, we combined the original unfiltered scRNA data for each patient’s AML sample as well as the healthy bone marrow sample. We normalized these samples together by median library size and applied a natural log transformation. It was necessary to return to the unfiltered data for this task since the highly variable gene filter was applied before Decipher removed some genes of interest. We then subsetted the cells to match those that were run through Decipher previously and plotted the expression of the selected genes (from the unfiltered set) in the previously obtained Decipher space. *Scanpy* plotting was utilized, with the colorbar range set to the .02-.09 percentiles of the data. For the scATAC-seq data, we created an AnnData object from the smoothed gene scores as described above and plotted using the projected Decipher coordinates (**Fig. SI3**).

### Motif Analysis in scATAC-seq

To obtain the set of TFs for motif analysis, we examined the time of maximum expression in each of the top 50 disrupted TFs (according to the combined disruption metric). If the time of maximum expression was before the estimated time of NPM1 mutation (e.g., at pseudotime 50 for AML1), we designated the TF to be “early peaking.” We then investigated whether the early peaking TFs could be putative regulators of other disrupted TFs, explaining the propagation of dysregulated mechanisms leading to the altered transcriptional landscape (**Fig. 6b,c**). Potential targets were thus defined as the set of remaining TFs (peaking after NPM1 mutation). Motif searching was performed using motifmatchR [105], with the JASPAR2020 [106] database of motifs and the Hg38 genome.

For each patient, we identified clusters obtained from the ArchR Iterative LSI analysis that corresponded to either blast 0 (AML1,3) or blast 1 (AML2). The ArchR reproducible peak set for that cluster, containing pseudo-bulked peak data, was used as the input to motifmatchR. To transform the result into a matrix of motifs by targets, we grouped motif scores by the target’s nearest gene and calculated the mean score for all scores belonging to the same gene.

To evaluate the significance of motif enrichment, we defined a background (null) set of TFs as follows: for each patient scRNA-seq data, we filtered out genes expressed in fewer than 1 percent of cells. This threshold was selected based on the proportion of PROM1-expressing cells in the patient samples, e.g., 125 out of 13834 cells in AML3, since we expected PROM1 to be a good example of a gene expressed in the rare immature cells of interest.

We then limited the null set to TFs that were not among the target set. P-values were computed by bootstrapping: we sampled the null set 10,000 times for a subset of TFs that were the same size as the target set. We then computed the mean motif score for all TFs in the null set to create the null distribution. The p-value was obtained by counting the number of samples in the null set whose score was greater than the mean motif score of our target set of TFs. We reported the p-values for the TFs that were significantly enriched in either all three patients (in which case the maximum p-value was reported) or significantly enriched in two out of three patients (in which case the maximum among the two significant p-values was reported).

In all three patients, we found significant motif enrichment in HOX TFs, including HOXB4 (motif JASPAR ID MA1499.1, p < 0.0458) and HOXB6 (MA1500.1, p < 0.032) and FOSL1 (MA0477.1, p < 0.0057; MA0477.2 0.0086), suggesting their putative role in regulating TFs driving AML transformation (\textbf{Fig. 6d}).

In two out of three patients, we found significant motif enrichments in GATA2 (MA0036.1, p < 0.0115; MA0036.2, p < 0.0423; MA0036.3, p < 0.0309), MYC (MA0147.3, p < 0.045), NR4A2 (MA0160.1, p < 0.0272), MAFF (MA0495.1, p < 0.0365), ATF3 (MA0605.2, p < 0.0316), NFKB2 (MA0778.1, p < 0.0292), RELB (MA1117.1, p < 0.0109), KLF2 (MA1515.1, p < 0.0468), ZNF274 (MA1592.1, p < 0.009) (**Fig. SI3**).

## Footnotes

1 Because the Gaussian Process is used here only to define the distribution of the observations, and not to sample an unobserved latent variable, there is no computational difficulty in using it.

